# Co-transcriptional RNA processing boosts zygotic gene activation

**DOI:** 10.1101/2024.09.14.613088

**Authors:** Jingzhao Xu, XiaoJing Li, Xiaowen Hao, Xinyun Hu, Shaoqian Ma, Yantao Hong, Jing Zhang, Dingfei Yan, Haiteng Deng, Jie Na, Xiong Ji, Zai Chang, Xiaohua Shen

**Affiliations:** School of Basic Medicine, Tsinghua University, Beijing 100084, China; School of Life Sciences, Tsinghua University, Beijing 100084, China; Key Laboratory of Cell Proliferation and Differentiation of the Ministry of Education, School of Life Sciences, Peking-Tsinghua Center for Life Sciences, Peking University, Beijing 100871, China

**Keywords:** Pol II CTD, co-transcriptional RNA processing, non-coding genome, zygotic gene activition, totipotency

## Abstract

Transcription decodes protein-coding genes and interprets regulatory information embedded in the genome by generating RNA. In eukaryotes, gene transcription is coupled with RNA processing via the carboxyl terminal domain (CTD) of RNA polymerase (Pol) II, which enhances messenger RNA (mRNA) production. We propose that co-transcriptional RNA processing is essential for zygotic gene activation (ZGA), transitioning the transcription program from noncoding to protein-coding after fertilization. Truncating the CTD in mouse cells disrupts this coupling, halting global mRNA synthesis and increasing noncoding RNA (ncRNA) levels through enhanced intergenic transcription and RNA stabilization. CTD truncation also triggers epigenetic reprogramming and nuclear reorganization towards totipotency, resembling early cleavage embryos. Mechanistically, the CTD restrains nonproductive polymerase activity in noncoding sequences, while at protein-coding genes requiring RNA processing, it promotes elongation by facilitating polymerase promoter-proximal pausing, transcription directionality, and velocity. Longer CTD lengths enhance gene activity, likely evolving to accommodate the increasing noncoding sequences in mammalian genomes.

## Introduction

The central dogma of molecular biology outlines genetic information flow from genes to mRNA to proteins^1^. However, about 98% of mammalian genomes consist of noncoding sequences that can be transcribed but do not encode proteins^2–5^. Understanding these noncoding regions is crucial for fully comprehending the genome. Transcription is the process by which genomic information is decoded into RNA molecules. Eukaryotic gene transcription is intricately linked with RNA processing, mediated by the CTD of the largest subunit (RPB1) of RNA Pol II^6–16^. The CTD plays a critical role in facilitating efficient RNA processing; truncation of the CTD inhibits capping, splicing, and cleavage-polyadenylation of mRNA precursors^15–22^. While this coupling enhances mRNA production, its broader biological significance remains unclear.

Transcription begins post-fertilization in a conserved process known as zygotic gene activation (ZGA) across animals and plants^23–31^. In mice, ZGA occurs in three distinct stages. Initially, genome-wide non-productive transcription is observed in one-cell embryos (pre-ZGA) ^32–34^. Subsequently, a minor wave of transcription involves about one hundred genes in the early two-cell stage (minor ZGA), followed by a major wave where several thousand genes are highly activated (major ZGA). Minor ZGA must precede major ZGA for a successful maternal-to-zygotic transition and embryo development beyond the two-cell stage^35,36^. Despite its critical role in initiating embryonic development across metazoans, the fundamental principles underlying ZGA remain elusive.

Mechanisms like maternal regulators, chromatin-mediated processes, and the DNA-to-cytoplasm ratio or cell-cycle duration partially explain genome activation timing but have limited cross-species applicability^37–45^. Disruption of maternal transcription factors^44,46–52^, RNA-binding proteins (RBPs) ^45,53^, and chromatin modifiers^54–56^ can impair ZGA onset, but doesn’t fully explain the shift from global transcription to gene-specific activation. *NANOG*, *SOXB1*, and *POU5F3* are critical for ZGA in zebrafish but expressed post-ZGA in *Xenopus* and mammals^57,58^. Species-specific patterns also exist in global DNA methylation and histone modifications during ZGA^59^. Rapidly developing organisms like flies and mosquitoes exhibit short early zygotic genes linked to unique cell-cycle dynamics absent in slower-developing animals^24,26,27,29^.

Interestingly, maternal transcripts, which are typically long, evolutionarily older, and conserved across species, are enriched in housekeeping functions such as RNA processing and translation^60^. These functions suggest their roles in early embryogenesis through RNA-driven mechanisms. In contrast, early zygotic genes are short, evolutionarily younger, and species-specific^60–64^. For instance, only 40% of human ZGA genes are shared with mice, and just 8.5% are common across humans, mice, and cattle^41^. This discrepancy aligns with the flexible ZGA period influenced by species-specific mechanisms, shifting from an initially less conserved to a highly conserved phase of basic body axis formation, as described by the developmental hourglass model^60,61,64–66^. These observations suggest that conserved processes, rather than species-specific factors, likely govern global gene activation during ZGA.

## Results

### Inhibiting RNA splicing blocks ZGA in mouse embryos

Indeed, despite species-specific differences in minor-ZGA (totipotency) genes, they commonly have shorter lengths and fewer introns compared to maternal transcripts, major-ZGA genes, and pluripotency genes expressed in the inner cell mass (ICM) of blastocysts and their derivative embryonic stem cells (ESCs) (Figure 1A). Using 5-ethynyl uridine (EU) labeling to track ongoing transcription, we detected nascent RNA in mid-one-cell embryos at the PN3 stage (pre-ZGA), with progressively stronger signals in late one-cell (PN5) and late two-cell embryos (Figures 1B and S1A). This suggests that early embryonic transcription, likely producing ncRNA, occurs before major ZGA. Unlike ncRNA production, mRNA synthesis requires co-transcriptional processing, involving 5’ capping, intron splicing, 3’ cleavage and polyadenylation, RNA modification, packaging, and nuclear export or RNA decay^14,67–69^. To test a role of co-transcriptional RNA processing in ZGA, we began by inhibiting RNA splicing through targeting small nuclear RNA (snRNA) or ribonucleoprotein complexes (snRNPs) critical for splicing junction recognition^70^. This involved microinjection of antisense morpholino oligonucleotides (AMOs) for U1/U2 snRNA and the use of Pladienolide B (PlaB), which targets the SF3B complex of U2 snRNP^71^. Splicing inhibition led to developmental arrest in over 95% of embryos at the two-cell stage, while control embryos progressed to the morula or blastocyst stages (Figures 1C and 1D). RNA-seq analysis showed that it blocked major ZGA but not minor ZGA, while inhibiting transcription with 5,6-Dichloro-1-β-D-ribofuranosylbenzimidazole (DRB) blocked both (Figures 1E-G and S1B-D). In addition, PlaB-treated embryos sustained aberrant high-level expression of minor-ZGA genes, impeding the move from the two-cell program to later stages (Figures 1G and S1C). Thus, RNA splicing appears crucial for transiting from minor to major ZGA. Consistently, inhibition of RNA splicing in mouse and human ESCs, either chemically or by knockdown of spliceosomal components, has been reported to induce totipotent reprogramming with ZGA-like gene signatures^72–74^. However, the underlying mechanism for this phenomenon remains obscure.

**Figure 1.**
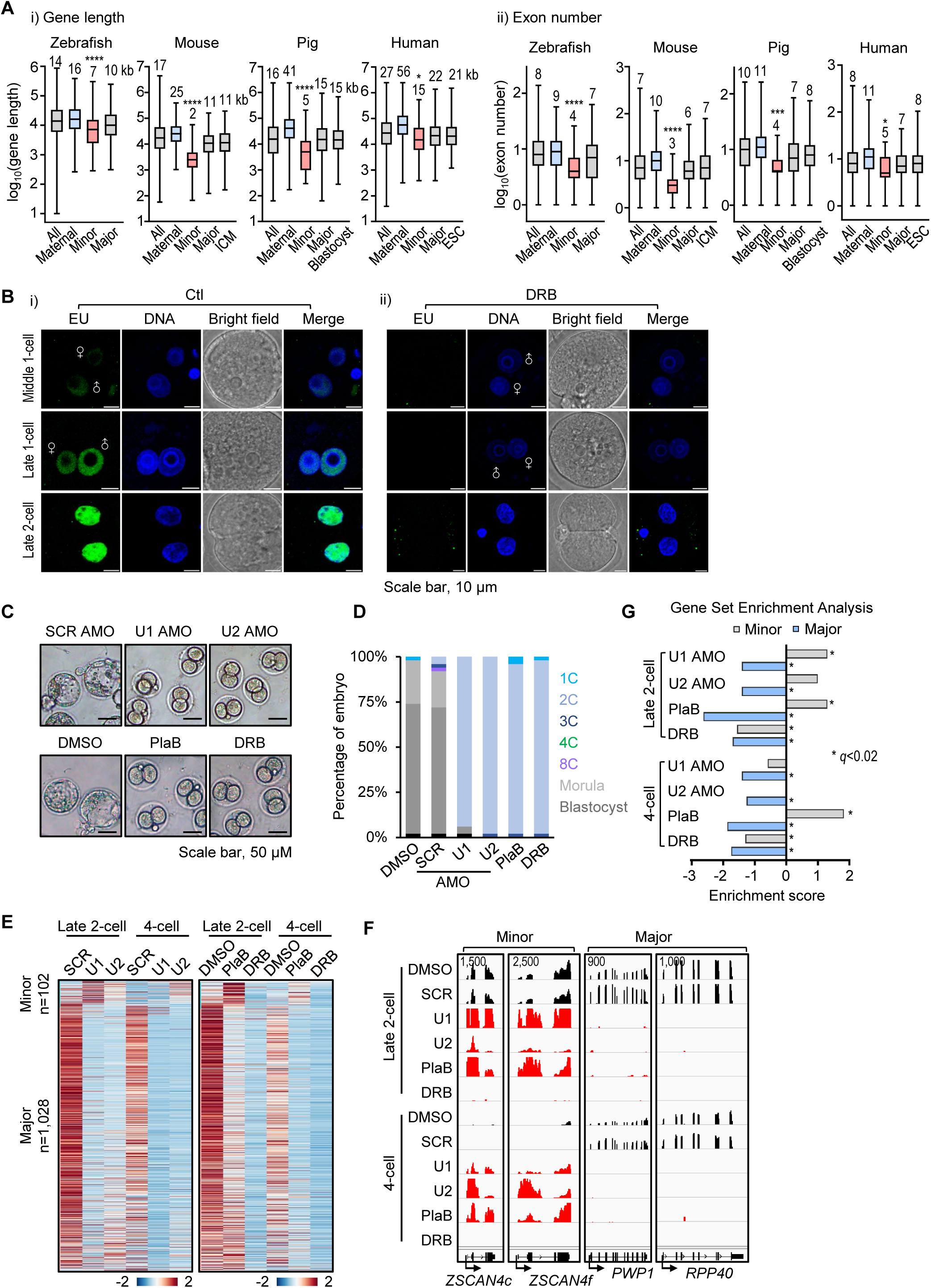
Inhibition of RNA splicing blocks ZGA. **(A)** Box plots illustrate that minor-ZGA genes have shorter lengths and fewer introns compared to all annotated protein-coding genes, maternal genes, and genes expressed in major-ZGA stage, blastocysts, or ESCs across the indicated species. Box plots display the 5th, 25th, 50th, 75th, and 95th percentiles, with the median value indicated. Statistical significance was determined using a two-tailed Mann-Whitney-Wilcoxon test, with *P*-values categorized as follows: > 0.1234 (ns), < 0.0332 (*), < 0.0021 (**), < 0.0002 (***), < 0.0001 (****). The numbers of analyzed genes in the gene-length (i) and exon-number (ii) analyses are shown before and after the slash, respectively. Zebrafish: all (*n =* 25,432/14,692), maternal (*n =* 1,243/879), minor-ZGA (*n =* 233/133), major-ZGA (*n =* 523/265); Mouse: all (*n =* 22,258/19,561), maternal (*n =* 1,009/945), minor-ZGA (*n =* 103/51), major-ZGA (*n =* 1,030/917), ICM (*n =* 528/465); Pig: all (*n =* 22,040/14,472), maternal (*n =* 3,023/2,507), minor-ZGA (*n =* 74/33), major-ZGA (*n =* 1,228/917), blastocyst (*n =* 1,331/1,108); Human: all (*n =* 20,030/5,426), maternal (*n =* 2,825/827), minor-ZGA (*n =* 132/34), major-ZGA (*n =* 914/268), ESC (*n =* 1,341/375). Refer to Methods and Table S1. **(B)** Representative images of early mouse embryos labeled with 5-ethnyl-uridine. EU-labeled RNA: green; DNA: DAPI, blue. Control (Ctl): DMSO. DRB serves as the negative control. Scale bar, 10 μm. **(C-D)** Inhibiting RNA splicing resulted in embryonic arrest at the two-cell (2C) stage. Panel (**C**) shows representative embryos, and panel (**D**) shows the quantification of embryos at various developmental stages. Controls: scramble (SCR) AMO and mock (DMSO) treated embryos; splicing-inhibited: U1 AMO, U2 AMO, and PlaB; and transcription-inhibited: DRB. Data are derived from two biological replicates. Scale bar, 50 μm. **(E)** Heatmap visualization of minor and major ZGA genes in embryos collected at the late two-cell (48 hphCG) and four-cell stages (60 hphCG). Inhibition of splicing blocked major ZGA but not minor ZGA, while inhibiting transcription blocked both processes. Expression levels of gene exons were normalized using spike-in RNA, which was added into an equal number of embryos and averaged across three biological replicates (refer to Methods and Table S2). The number of ZGA genes analyzed (FPKM > 0): minor, *n =* 102, and major *n =* 1,028. The color scale corresponds to z-score normalized FPKM. **(F)** Integrative Genomics Viewer (IGV) showing RNA-seq signals of representative minor and major ZGA genes, with the signal scale shown at the top left. **(G)** Gene Set Enrichment Analysis (GSEA) of minor- and major-ZGA genes. Embryos with splicing inhibition (U1 or U2 AMO or PlaB) or transcription inhibition (DRB) showed downregulation of major-ZGA genes compared to control embryos collected at both late two-cell and four-cell stages, while minor-ZGA genes were upregulated in PlaB-treated embryos at both stages. The x-axis denotes the Normalized Enrichment Score (NES). Significant enrichment (q < 0.02) is indicated by an asterisk. For additional details, see also Figure S1C.

### CTD truncation induces a totipotency gene program

It is well documented that truncating the CTD in yeast and mammalian cells inhibits pre-mRNA capping, splicing, cleavage, polyadenylation, and mRNA nuclear exporting^15–22^. The CTD forms a large unstructured domain that serves as a binding platform for RNA processing machineries^7,13,15,16,75^. It contains repeated heptapeptide motifs, ranging from 5 repeats in *P. yoelii* to 26 in *S. cerevisiae*, and up to 52 repeats in mammals^76,77^. The CTD regulates transcription initiation and elongation through phosphorylation at the serine 5 and 2 sites (S5P and S2P) of the heptapeptide^7,13,75^. Given its role in coupling transcription and pre-mRNA processing, we sought to disrupt this coupling by truncating the CTD from the Pol II and investigate its impact on transcription activity across the genome.

We utilized a mouse ESC line engineered with an auxin-inducible degron (AID) tag fused to the carboxyl terminus of endogenous RPB1^78^. Treatment with the auxin analog indole-3-acetic acid (IAA) led to CTD truncation while preserving the catalytic amino-terminal domain (NTD) of RPB1, which maintains a compact structure with other Pol II subunits (Figures 2A and S2A). This truncation abolished CTD phosphorylation (S5P and S2P) (Figure 2A). Mass spectrometry analysis of insoluble nuclear fractions, including the chromatin and ribonucleoprotein (RNP) mesh, revealed a significant decrease in 577 proteins involved in mRNA processing, splicing, and transport, transcription, chromatin organization, and the cell cycle 24 hours post-IAA treatment (Figures S2B, C). This aligns with the CTD’s role as a central hub for interactions with RNA processing and transcription machineries^13,15,16,67,79,80^.

**Figure 2.**
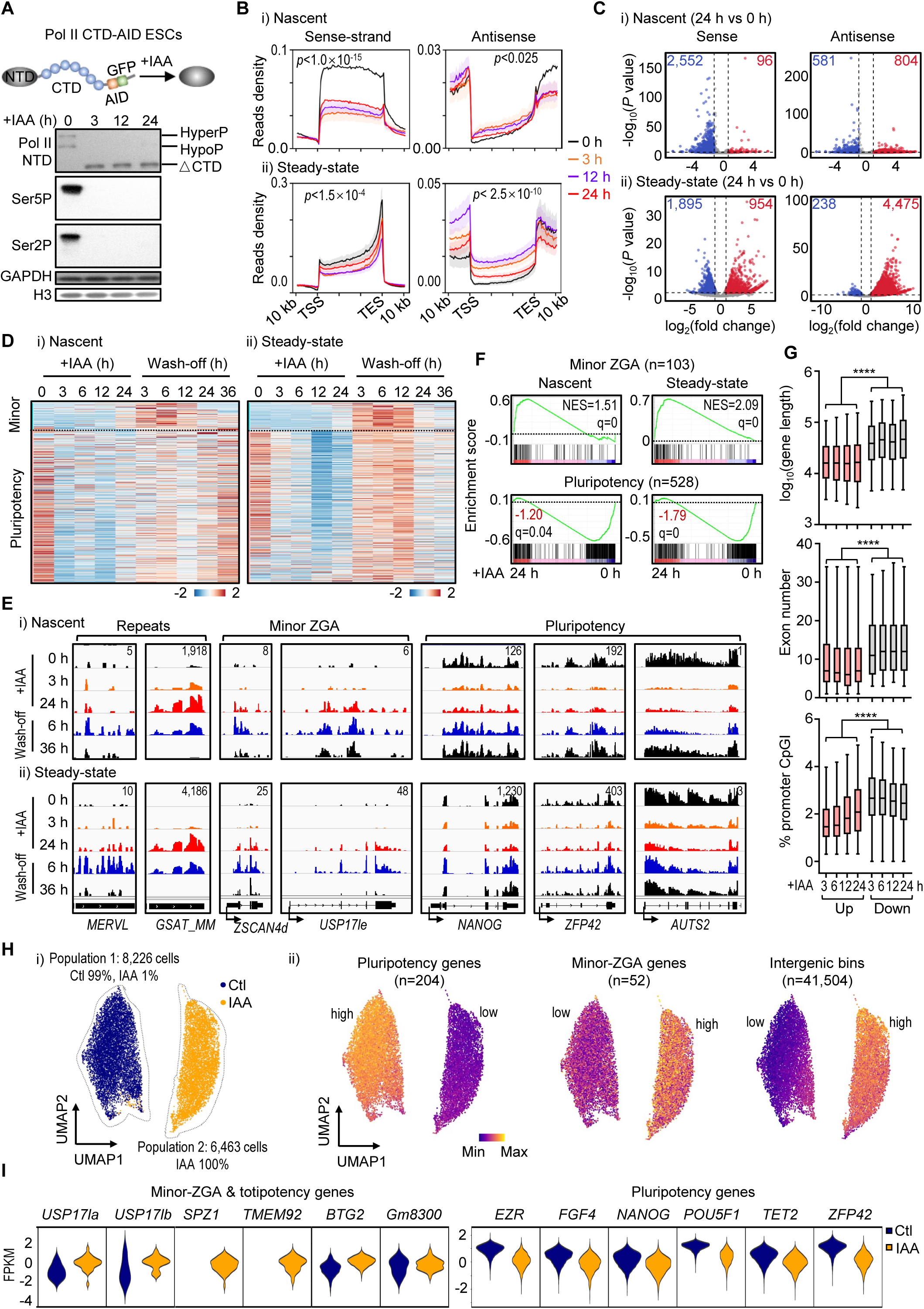
Truncation of the Pol II CTD downregulates global gene transcription but upregulates minor-ZGA genes. **(A)** Pol II CTD-AID ESCs. The upper diagram shows the endogenous RPB1 tagged with AID- and GFP-tag at its carboxyl terminus. Upon the addition of IAA, the CTD is truncated from the compact core structure containing the amino-terminal domain (NTD) of RPB1 of the Pol II. The western blots below show the rapid truncation of the CTD and the loss of phosphorylation after IAA treatment at the indicated time points, with antibodies listed on the left. Protein bands corresponding to hyperphosphorylated (hyperP), hypophosphorylated (hypoP), and CTD-truncated (ΔCTD) Pol II are indicated. GAPDH and histone H3 are included as loading controls. **(B)** Metagene analysis of (i) nascent RNA by EU-seq and (ii) steady-state RNA by rRNA-depletion RNA-seq in Pol II CTD-AID ESCs after the addition of IAA treatment for 3 to 24 hours, as indicated. RNA read signals in the sense (left) and antisense (right) strands of all protein-coding genes (*n =* 22,258) are shown. Read density was normalized to cell numbers using spike-in controls. Data represent mean values from two biological replicates, with shadings representing the standard error of the mean (SEM). Significance was tested with a two-sided Kolmogorov-Smirnov test, with *P*-values as indicated. **(C)** Volcano plots showing differential expression of genes in the sense strand (left) and antisense strand (right) post 24-hour IAA treatment compared to untreated control (0-hour): (i) Nascent RNA (sense, *n =* 8,429 genes with sum counts > 5; antisense, *n =* 6,223 genes with sum counts > 20) and (ii) Steady-state RNA (sense, *n =* 14,759 genes with sum counts > 5; antisense, *n =* 11,967 genes with sum counts > 20). Genes with log_2_(fold change) > 1 and *P* < 0.05 are highlighted in blue (downregulated) and red (upregulated), with gene numbers annotated above. *P*-values were obtained from a two-tailed t-test based on two biological replicates. **(D)** Heatmaps displaying relative gene expression during the time course of adding IAA for 24 hours followed by a 36-hour wash-off: (i) nascent and (ii) steady-state RNA for minor-ZGA (*n =* 92, FPKM > 0) and pluripotency genes (*n =* 523, FPKM > 0). Expression levels of gene exons were normalized by spike-in RNA added into the same number of cells and averaged from two biological replicates for IAA treatment and three replicates for wash-off samples. The color scale corresponds to z-score normalized FPKM. **(E)** IGV snapshots of nascent (i) and steady-state RNA (ii) for selected genomic loci in Pol II CTD-AID ESCs following IAA treatment and wash-off at specified time points. Transcription direction (bottom) and the signal scale (top right) for grouped tracks are provided. The signals represent averages from two biological replicates for IAA treatment and three replicates for wash-off samples, normalized using spike-in controls. **(F)** GSEA of minor-ZGA (top) and pluripotency genes (bottom) in Pol II CTD-AID ESCs treated with IAA for 24 hours. Gene counts, NES, and false discovery rate (FDR) q-values are as indicated. **(G)** Box plots illustrate that upregulated genes are significantly shorter in length, and have fewer exons and lower CpG-island density at their promoters compared to downregulated genes. This analysis was based on steady-state gene expression at various time points adding IAA. The numbers of analyzed genes in the gene-length (i) and exon-number (ii) and promoter CpG density (iii) analyses are shown sequentially before and after the slash, respectively. Upregulated: 3-h = 342/307/305, 6-h = 669/592/599, 12-h = 883/793/797, 24-h = 954/874/872. Downregulated: 3-h = 1,531/1,453/1,441, 6-h = 2,832/2,652/2,639, 12-h = 4,056/3,836/3,806, 24-h = 1,895/1,786/1,780. The box plots display the 5th, 25th, 50th, 75th, and 95th percentiles, excluding outliers. Statistical significance was determined using a two-tailed Mann-Whitney-Wilcoxon test, with *P*-values indicated as follows: < 0.0001 (****). For further details, refer to Table S4. **(H-I)** Single-cell RNA-seq analysis of CTD-AID ESCs treated with or without IAA for 24 hours. In panel (**H**), the UMAP plot reveals two principal clusters: one mainly composed of control cells without IAA treatment (*n =* 8,140 cells) and another comprising IAA-treated cells (*n =* 6,549), with only 86 IAA-treated cells falling into the control cluster. The corresponding expression levels of minor-ZGA genes (*n =* 52), pluripotency genes (*n =* 204), and intergenic bins (*n =* 41,504) are depicted in the UMAP plots. Scale bars represent gene expression levels. In panel (**I),** violin plots show the expression distribution of representative minor-ZGA (totipotency) and pluripotency genes in individual control and IAA-treated cells.

We performed sequencing analysis of EU-labeled nascent RNA and total cellular RNA depleted of ribosomal rRNA. Acute CTD truncation led to a global decrease in gene-body sense-strand transcripts, and an increase in antisense RNA within the gene-body and upstream promoter regions, observed in both nascent and steady-state RNA levels (Figures 2B, C and S2D). Starting at 3 hours, there was a more drastic reduction in nascent RNA levels compared to steady-state mRNA, with 3,334 genes downregulated and only 54 genes showing significant upregulation (Figures 2C and S2D). This pattern persisting until 24 hours indicates a global blockade of mRNA synthesis. Particularly, pluripotency genes (*n =* 528) were rapidly downregulated at 3 hours (Figures 2D, E), followed by upregulation of minor-ZGA genes (*n =* 103, such as *ZSCAN4* and *USP17* families). *MERVL* and major satellite repeats also increased 24 hours post-IAA treatment (Figures 2D-F and S2E). Upregulated genes, characteristic of minor-ZGA genes, notably exhibit shorter lengths, fewer introns, and lower CpG-island density at the promoter compared to downregulated genes (Figures 1A, 2G and S2F, G). Removal of IAA gradually restored pluripotency genes to normal levels, while minor-ZGA genes initially increased but eventually decreased after 24-hour IAA wash-off (Figures 2D, E), underscoring the CTD’s role in complete activation of the gene program. Single-cell RNA-seq analysis of 24-hour IAA-treated cells revealed a profound divergence from control ESCs, with nearly all treated cells displaying elevated totipotency and reduced pluripotency gene programs (Figures 2H, I).

### Genome-wide pervasive transcription in CTD-truncated cells and minor-ZGA embryos

In pre- and minor-ZGA stages, we observed a striking common feature: transcription is pervasive and indiscriminate across the entire genome in multiple species. This was demonstrated by our analysis of eight published embryonic transcriptomes, detecting nascent, total or poly(A) RNA in bulk or single cells in mouse^32,55,81–83^, human^44,84^, and pig^85^. Specifically, in mouse, to quantify non-gene transcription, we divided intergenic regions into 35,553 proximal (< 10-kb from genes) and 11,418 distal bins (> 10-kb from genes) (Figure 3A), with distal bins presumably less influenced by genes compared to proximal bins. The proximal and distal bins have median lengths of 10-kb and 35-kb, respectively, representing 11.5% and 46% of the genome. In comparison, 24,155 genic bins have a median length of 16-kb, constituting 42.5% of the genome (Figures 3A).

**Figure 3.**
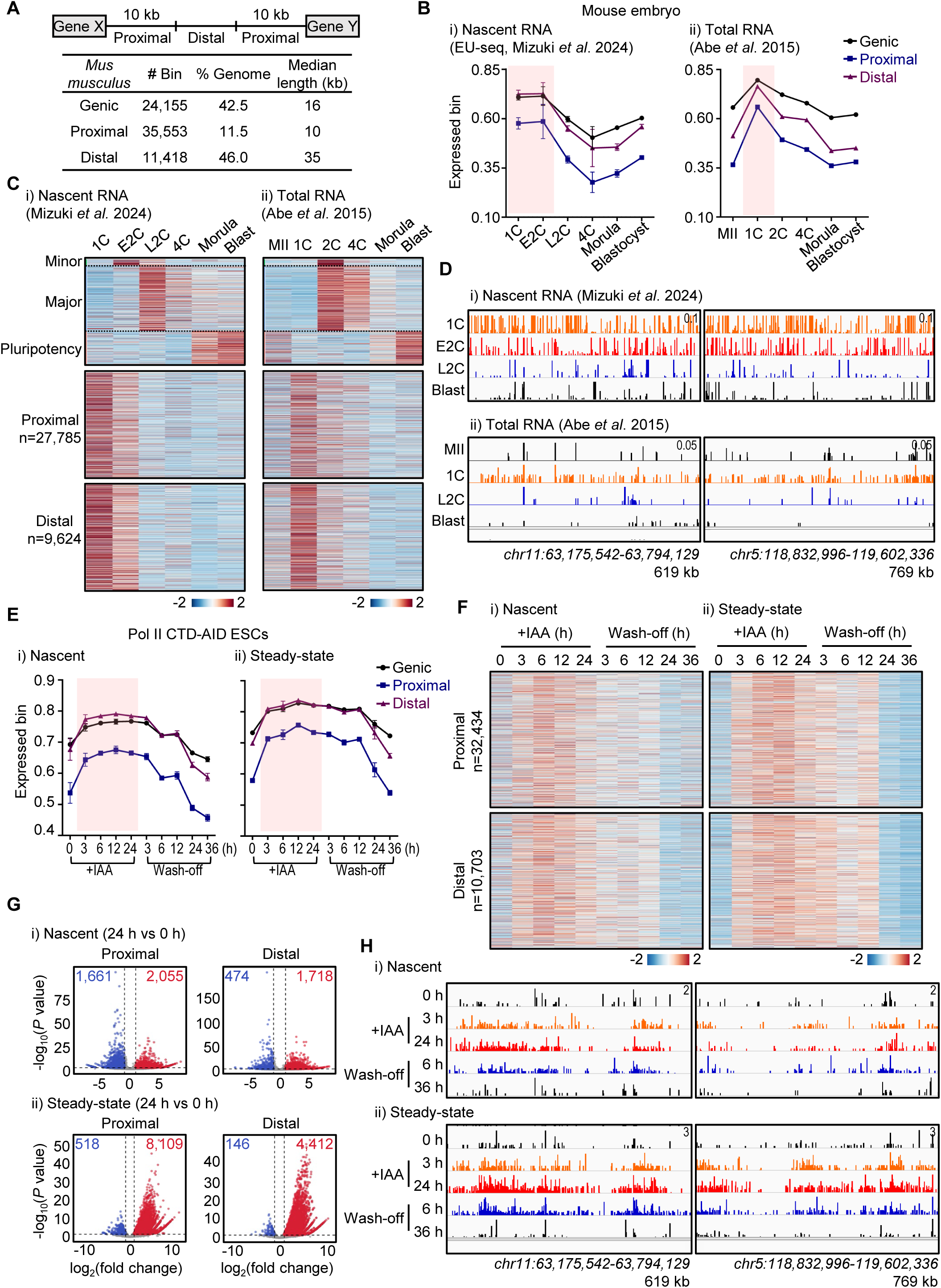
Pervasive intergenic transcription in CTD-truncated cells and early embryos. **(A)** Classification of the genome into genic and intergenic bins. Proximal and distal intergenic bins are defined by their distance to a gene, within 10 kb and beyond 10 kb, respectively. The bin number, percentage of genome coverage, and median length are indicated. **(B-D)** Analysis of nascent and steady-state RNA in early embryos based on published EU-seq and total RNA-seq. Panel (**B**) shows the percentages of transcribed bins (FPKM > 0) during early embryonic development. The pre- and minor-ZGA stages, highlighted in pink shading, exhibit the highest level of expressed genic and intergenic bins in both nascent and steady-state RNA levels. This suggests a greater proportion of indiscriminate genome-wide transcription compared to other embryonic stages. Panel (**C**) shows a heatmap illustrating the shift from noncoding to coding transcription during the transition from minor to major ZGA. Numbers of analyzed genes and intergenic bins: 97 minor-ZGA genes; 1,011 major-ZGA genes ; 514 pluripotency genes; 27,785 proximal and 9,624 distal bins. The FPKM values represent the mean of two biological replicates for EU-seq, with a filtering threshold of minimum FPKM > 0. Panel (**D**) shows IGV snapshots of EU-seq (top) and total RNA-seq (bottom) for representative intergenic loci at different stages of embryo development. Genome coordinates and scale are provided. **(E-H)** Analysis of nascent (EU-seq) and steady-state RNA levels in CTD-AID cells treated with IAA for 24 hours followed by a 36-hour wash-off of IAA. Panel (**E**) shows increasing percentages of transcribed genic and intergenic bins (FPKM > 0) in both nascent and steady-state levels during IAA treatment, highlighted in pink shading. Data points represent means from two biological replicates for the IAA treatment and three replicates for the wash-off period; error bars are SEM. In panel (**F**), heatmaps illustrate relative expression changes in proximal and distal intergenic bins, in nascent (left) and steady-state levels (right). The ‘*n*’ values denote the number of intergenic bins with minimum FPKM > 0. Expression levels are averaged from two biological replicates for IAA treatment and three replicates for wash-off. In panel (**G),** volcano plots display differential expression of intergenic bins in CTD-AID ESCs following 24-hour IAA treatment versus the untreated control (0-hour). Results from (i) nascent EU-seq for proximal (left, *n =* 15,405) and distal (right, *n =* 6,895) intergenic bins, as well as (ii) steady-state rRNA-depletion RNA-seq for proximal (left, *n =* 19,030) and distal (right, *n =* 7,577) intergenic bins are shown, with the criteria of summed read counts > 20 in each individual analysis. Bins with log_2_(fold change) > 1 and *P* < 0.05 are highlighted in blue (downregulated) and red (upregulated), with the number of significantly changed bins annotated above each plot. Normalization was done using spike-in controls added to an equal number of cells. *P*-values are calculated using a two-tailed t-test from two biological replicates. In panel (**H**), IGV snapshots show nascent (i) and steady-state (ii) RNA levels for representative intergenic loci in CTD-AID ESCs during IAA treatment and after wash-off at the indicated time points. Genome coordinates and scale are provided.

Summing RNA read counts in each bin revealed that one-cell (1C, pre-ZGA) to early two-cell embryos (E2C, minor ZGA) exhibit the highest percentages of genic and intergenic bins being transcribed, compared to late two-cell (L2C, major ZGA), four-cell (4C), morula, and blastocyst embryos (Figures 3B and S3A). Heatmaps and genome tracks also revealed widespread RNA reads in proximal and distal intergenic bins in 1C and E2C embryos, which significantly decreased at L2C, 4C, and blastocyst embryos (Figures 3C, D and S3A). This downregulation of intergenic RNA reads coincides with the activation of minor- and major-ZGA genes (Figures 3C and S3A). Similarly, in human embryos, a period of pervasive noncoding transcription at the two-to-four-cell stages is followed by a shift to coding gene-centered transcription at the eight-cell stage, coinciding with the onset of major ZGA (Figure S3B). The noncoding-to-coding transition during pre-to major ZGA was also observed in pigs (Figure S3C).

Interestingly, despite a global decrease in mRNA levels following CTD truncation, the proportion of transcribed genic bins increased (Figure 3E). Additionally, transcribed proximal and distal intergenic bins also increased, with RNA levels in both nascent and steady states rising rapidly within 3 hours of CTD truncation (Figures 3E-H and S3D). This phenomenon was reversible by washing off IAA after a 24-hour treatment (Figures 3E, F). Unsupervised hierarchical clustering revealed that ESCs treated with IAA for 3 to 24 hours clustered with 1C and E2C embryos, early wash-off cells (24-hour IAA plus 3-12-hour wash-off) clustered with L2C and 4C embryos, and late wash-off cells (24-hour IAA plus 24-to-36-hour wash-off) clustered with blastocysts and ESCs (Figure S3E). Thus, acute CTD truncation attenuated global mRNA synthesis while simultaneously upregulating genome-wide intergenic transcription and a handful of minor-ZGA genes. These changes suggest a dynamic transition in the transcriptome from pluripotency to totipotency reminiscent of embryos before the major-ZGA stage.

### Compromised RNA decay upon CTD truncation

Starting at the 3-hour IAA treatment, we observed a consistently higher increase in steady-state intergenic ncRNAs compared to their nascent transcripts, while nascent mRNAs decreased more than their steady-state levels (Figures 2B, C, 3G, H and S2D, S3D). This suggests a rapid inhibition of RNA decay after 3 hours of CTD truncation. Indeed, pulse-chase experiments revealed significantly extended RNA half-lives in cells treated with IAA for 24 hours: the median half-life of mRNAs increased from 88 to 191 minutes, introns from 22 to 62 minutes, proximal ncRNAs from 27 to 88 minutes, and distal ncRNAs from 39 to 106 minutes, along with various ncRNA subtypes (Figures 4A-D and S4A-D). In control ESCs, mRNA half-life appears to be negatively correlated with nascent transcription levels and positively correlated with GC dinucleotide content. These correlations were disrupted after CTD truncation (Figures 4E-F and S4E). Moreover, both up- and down-regulated genes showed similar increases in RNA half-life (Figures 4G), indicating an indiscriminate enhancement of RNA stability. In early cleavage embryos, maternal RNA decay becomes active at the time of major ZGA, ensuring proper maternal-to-zygotic transitions^25,86–88^. Thus, compromised RNA decay upon CTD truncation is consistent with totipotency reprogramming.

**Figure 4.**
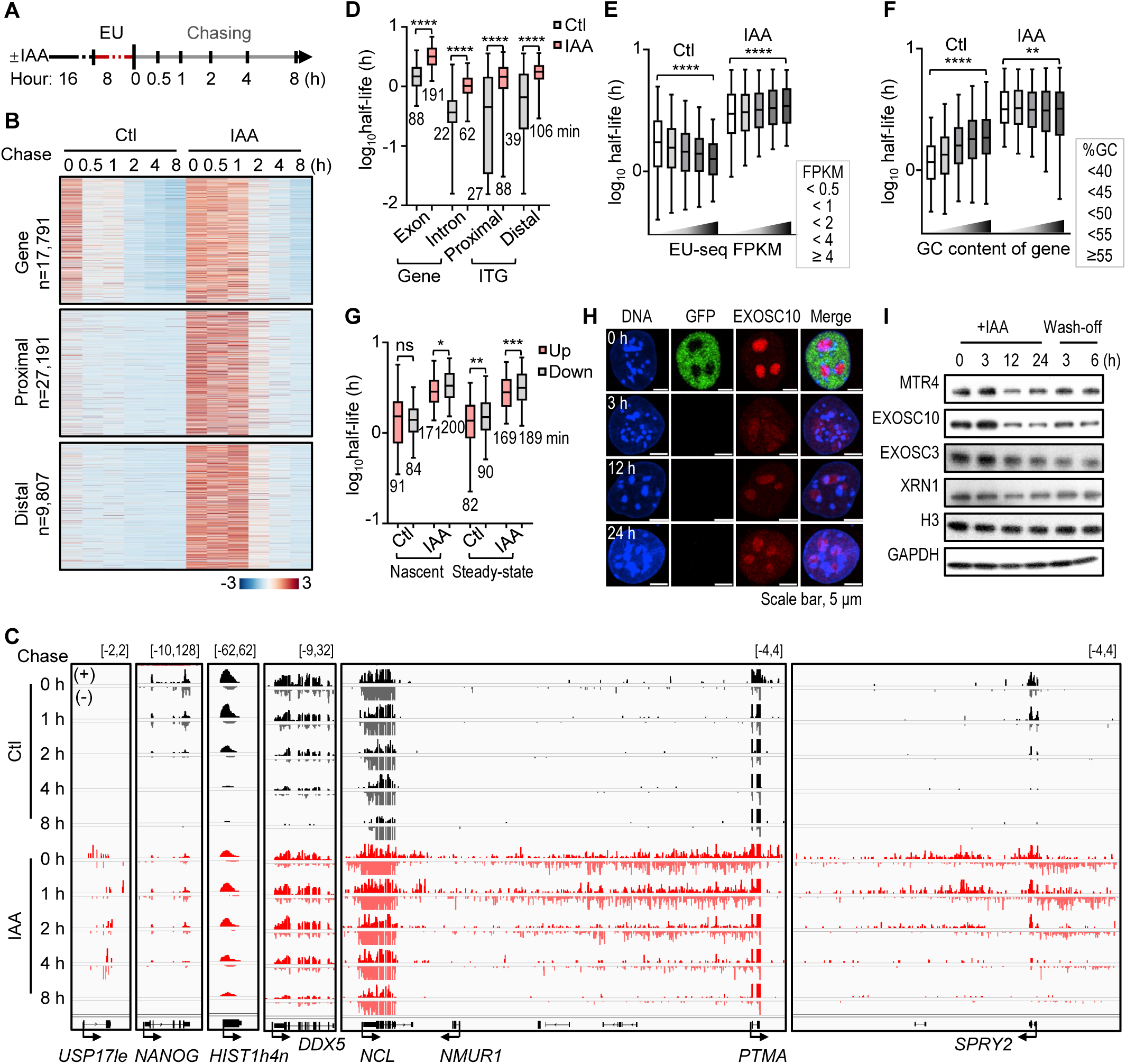
Compromised RNA decay upon Pol II CTD truncation. **(A)** Schematic workflow of pulse-chase experiments measuring RNA decay. Experimental group cells were treated with IAA for 16 hours prior to EU labeling for 8 hours. After removing EU, samples were collected at various chase time points for EU-seq analysis. IAA was continuously added throughout the entire process for the experimental group. **(B)** Heatmap showing EU-labeled RNA abundance of all protein-coding genes (*n =* 17,791, including exon and intron reads) and intergenic bins (proximal = 27,191, distal = 9,807) filtered with FPKM > 0 at indicated time points during the chase. Expression levels were averaged from two biological replicates using spike-in and z-score normalization. **(C)** IGV showing EU-labeled RNA signals at representative genes and intergenic regions across different time points during the chase, with strand orientation and scale indicated. **(D)** Box plots showing the half-life of mRNA genes (exon, *n =* 7,169; intron, *n =* 11,846), proximal (*n =* 12,204), and distal (*n =* 5,281) intergenic bins with and without IAA treatment. Only genes or regions with detectable RNA reads at 30 minutes or beyond can be used for half-life calculation, so fast-decaying transcripts within the 30-minute chase cannot be computed, potentially overestimating RNA half-lives. Nevertheless, all analyzed transcripts show enhanced stability. **(E)** Box plots showing the correlation between gene expression level and RNA stability. Gene categories were defined based on EU-seq FPKM of the control group (0-hour): FPKM < 0.5 (*n =* 1,634), < 1 (*n =* 1,392), < 2 (*n =* 1,702), < 4 (*n =* 1,296), ≥ 4 (*n =* 1,086). **(F)** Box plots showing the correlation between the GC content of genes and RNA stability. GC content categories: < 40% (*n =* 790), < 45% (*n =* 2,165), < 50% (*n =* 1,795), < 55% (*n =* 1,119), ≥ 55% (*n =* 365). In panels (**e**, **f**), the box plots display the 5th, 25th, 50th, 75th, and 95th percentiles, excluding outliers. *P*-values were calculated using an unpaired, one-way ANOVA Kruskal-Wallis test. **(G)** Box plot showing the mRNA half-life of upregulated (*n =* 44/198) and downregulated genes (*n =* 1,852/1,099) in nascent and steady-state RNA levels upon IAA treatment for 24 hours, with analyzed gene numbers shown before and after the slash, respectively. In panels (**d**, **g**), the box plots display the 5th, 25th, 50th, 75th, and 95th percentiles, with median values labeled. Statistical significance was determined using a two-sided paired Mann-Whitney-Wilcoxon test, with *P*-values denoted as follows: > 0.1234 (ns), < 0.0332 (*), < 0.0021 (**), < 0.0002 (***), < 0.0001 (****). **(H)** Immunofluorescence imaging in GFP-tagged CTD-AID cells treated with IAA over a 24-hour time period, showing EXOSC10 in red and DNA (DAPI) in blue. The loss of GFP indicates successful truncation of the CTD. Scale bar, 5 μm. **(I)** Western blot analysis of cell lysates from CTD-AID ESCs treated with IAA for 24 hours, followed by a 6-hour wash-off. Histone H3 and GAPDH were included as loading controls.

Within 3 hours of IAA treatment, the RNA exosome, responsible for 3’ to 5’ RNA decay^89–91^, exhibited reduced nuclear localization, particularly in the nucleolus, as shown by EXOSC10 immunostaining (Figures 4H and S4F). Western blots showed decreased levels of EXOSC10 protein and other exosome components (EXOSC3, MTR4) in 12 hours (Figure 4I). We speculate that in response to decreased global mRNA synthesis upon CTD truncation, cells upregulate both noncoding transcription and RNA stability to maintain overall RNA levels, underscoring the intricate link between RNA synthesis and degradation^91–93^. We previously reported that acute inhibition of the RNA exosome did not cause upregulation of minor-ZGA genes or intergenic transcription^93^, suggesting that the observed reprogramming effects are not solely due to compromised RNA decay.

### Altered Pol II binding mirrors embryonic transcription

Chromatin immunoprecipitation and sequencing (ChIP-seq) targeting the NTD of RPB1 revealed genome-wide changes in Pol II binding, consistent with alterations in nascent RNA (Figures 2B, 3F and 5A). Within 3 hours of CTD truncation, Pol II binding significantly increased in proximal and distal intergenic regions (Figure 5A), suggesting that the CTD normally restrains Pol II binding broadly across chromatin. In genic regions, Pol II binding diminished at the transcription start site (TSS) but increased along gene bodies and surrounding regions (Figures 5A-C and S5A), indicating abolished promoter pausing. In comparison, ZGA embryos exhibit atypical Pol II phosphorylation with nearly indistinguishable genic distributions of Ser2P and Ser5P, minimal promoter pausing, and a higher gene-body-to-promoter binding ratio (Figures S5A, B) ^36,94^. Thus, reminiscent of two-cell embryos the CTD-truncated Pol II binding profile, characterized by a lack of promoter pausing and increased gene-body and intergenic binding, contrasts with the classical Ser5P pattern at promoters and Ser2P along gene bodies observed in eight-cell embryos and ESCs.

**Figure 5.**
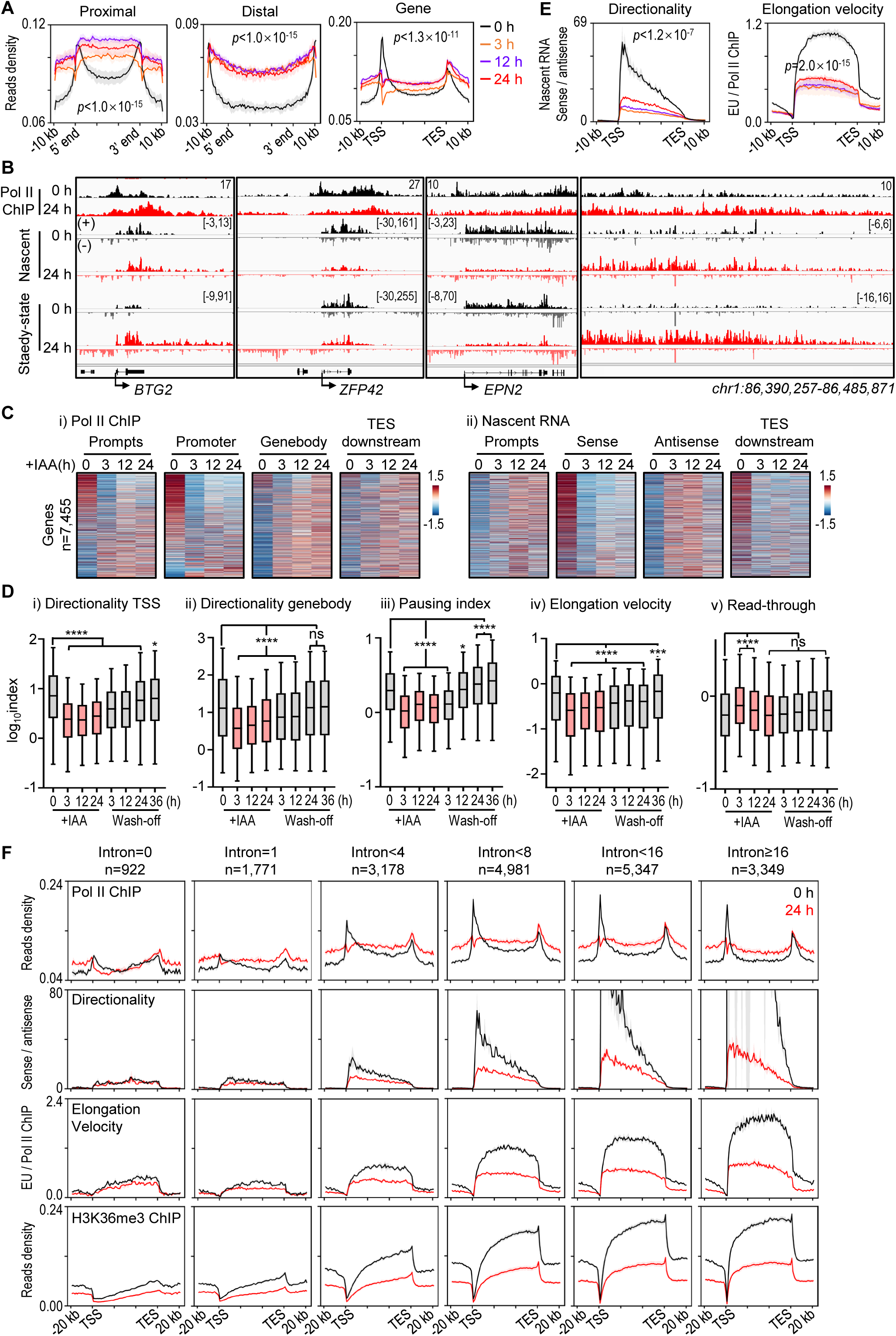
The Pol II CTD enhances transcription regulation and efficiency in complex genes. **(A)** Metaplot analysis of Pol II ChIP-seq targeting the NTD of RPB1. During a 24-hour IAA treatment, acute truncation of the Pol II CTD led to increased Pol II binding across proximal (*n =* 35,553) and distal (*n =* 11,418) intergenic regions. At protein-coding genes (*n =* 22,258), Pol II binding at the TSS was drastically reduced, while binding increased in the upstream, gene-body, and downstream regions. **(B)** IGV snapshots showing Pol II ChIP-seq, nascent EU-seq, and steady-state RNA-seq signals in representative genes and intergenic regions before and after a 24-hour IAA treatment. Strand orientation and scale are as indicated. **(C)** Heatmaps showing consistent Pol II binding (i, ChIP-seq) and nascent RNA levels (ii, EU-seq) during IAA treatments across various genomic regions. These include PROMPTs (2-kb upstream of gene TSS on the antisense strand), promoter (±300 bp around TSS on the sense strand), gene bodies (from 300 bp downstream of TSS to 300 bp upstream of TES on the sense strand), and TES-downstream regions (2-kb downstream of TES on the sense strand) for 7,455 genes showing detectable signals (maximum FPKM > 0) in each region. Each row represents different regions of the same gene. **(D)** Box plots illustrating the decreases in Pol II directionality around TSS (*n =* 2,224) and within gene bodies (*n =* 9,519), promoter pausing index (*n =* 5,724), and elongation velocity (*n =* 8,431), as well as an increase in read-through transcription beyond TES (*n =* 4,552) during a 24-hour IAA treatment. These changes returned to normal levels during a 36-hour wash-off period. Specific calculation methods are detailed in Figure S5A and Methods. The box plots display the 5th, 25th, 50th, 75th, and 95th percentiles, with outliers omitted. Statistical significance was determined using a two-tailed Mann-Whitney-Wilcoxon test, with *P*-values denoted as follows: > 0.1234 (ns), < 0.0332 (*), < 0.0021 (**), < 0.0002 (***), < 0.0001 (****). **(E)** Metaplots of transcription directionality and elongation velocity at protein-coding genes (*n =* 22,258) showing drastic reductions after acute CTD truncation following the addition of IAA for 3-24 hours. Transcription directionality was calculated as the ratio of EU-seq read density on the sense strand to the antisense strand. Elongation velocity of a gene was calculated as the ratio of EU-seq read density on the sense strand to Pol II ChIP read density. Significance was tested with a two-sided Kolmogorov-Smirnov test, with *P*-values as indicated. **(F)** Metaplots showing Pol II binding, directionality, elongation velocity, and the elongation histone mark H3K36me3 at distinct classes of genes categorized by exon numbers before and after a 24-hour IAA treatment. Also refer to Figure S5C. In panels (**A**, **E**, **F**), data represent mean values from three biological replicates for ChIP-seq and two biological replicates for EU-seq using spike-in normalization. Shadings represent SEM.

In addition, elongation velocity, measured by nascent sense-strand RNA signals normalized to Pol II ChIP density, markedly decreased (Figures 5D, E and S5A). Transcription directionality, assessed by the sense-to-antisense nascent RNA ratio at TSS and gene bodies, was compromised shortly after the IAA addition (Figures 5B, D, E and S5A). Conversely, transcription read-through, determined by RNA signals downstream of the transcription end site (TES) normalized to gene bodies, significantly increased (Figures 5B, D and S5A).

Positive correlations were observed between increasing intron numbers and promoter CpG-island densities with the degree of Pol II promoter pausing, transcription velocity, directionality, and the elongation histone mark H3K36me3 (Figures 5F and S5C, D). However, CTD truncation disrupted these correlations, particularly affecting genes with more introns and higher promoter CpG content (Figures 5F, and S5C, D). This indicates that the CTD enhances transcriptional regulation and efficiency for genes with complex structures. Correspondingly, genic transcription in E2C embryos exhibits lower directionality, velocity, and Pol II promoter pausing, along with higher read-through, compared to L2C or ICM stages (Figures 5B, E). We conjecture that in early embryos, the gradual establishment of co-transcriptional RNA processing accounts for the progressive increase in transcription regulation and efficiency, meeting the growing need for mRNA production of longer genes during development.

### Epigenetic and nuclear reprogramming towards early embryos

ChIP-seq revealed significant changes in chromatin states in CTD-AID cells after 24 hours of IAA treatment (Figures 6A-C and S6A). Gene activity-associated marks like H3K4me3, H3K36me3, and H3K27me3, decreased globally, while the repressive H3K9me3 mark notably increased across both genic and intergenic regions (Figures 6A, C and S6B). *<u>A</u>*ssay for *t*ransposase-*<u>a</u>*ccessible *<u>c</u>*hromatin using sequencing (ATAC-seq) revealed decreased TSS signals but increased gene-body and distal signals (Figures 6A and 6B). Notably, for all chromatin marks analyzed, peaks enriched in the ICM were generally lost in CTD-truncated cells, whereas gained peaks such as ATAC and broad H3K4me3 resembled those in two-cell embryos (Figures 6B). The drastic reduction of H3K36me3 in CTD-truncated cells aligns with the observation that embryonic H3K36me3 marks begin weakly in late two-cell embryos and strengthen by the eight-cell stage (Figures 6A, C and S6A, B) ^54,95^. The reduction in bivalent marks also corresponds with their transient nature in two-cell embryos, typically stabilizing only by the blastocyst stage (Figures 6A, C) ^96^. Additionally, positive correlations were observed between intron numbers and the enrichment of H3K4me3 and ATAC signals, as well as the depletion of H3K9me3 at the TSS (Figure 6C), indicating a genome-encoded regulatory mechanism for genes.

**Figure 6.**
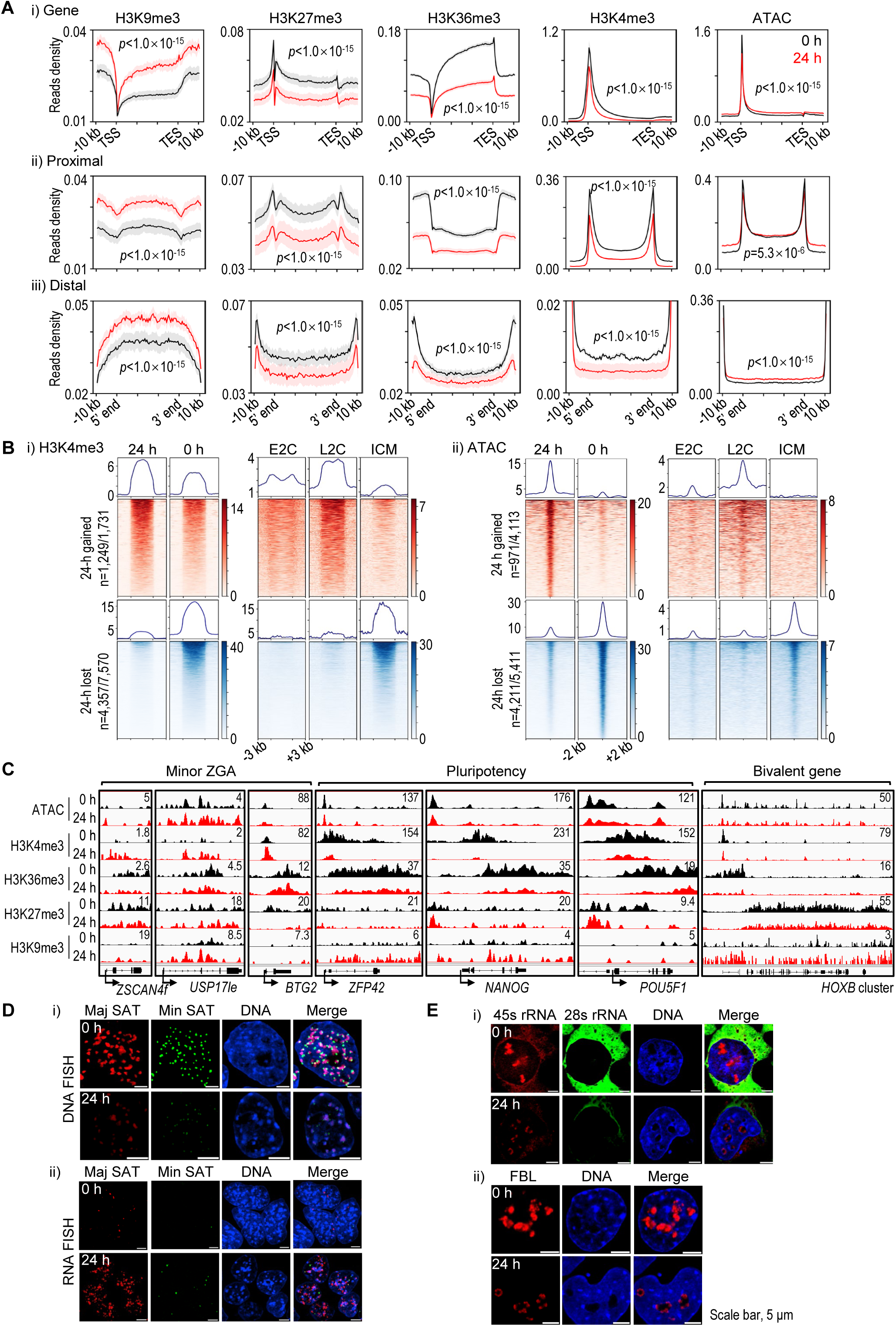
CTD truncation triggers epigenetic reprogramming and nuclear reorganization towards early embryos. **(A)** Analysis of histone modifications by ChIP-seq and chromatin openness by ATAC-seq before (0 h) and after a 24-hour (24 h) IAA treatment in CTD-AID ESCs. Panel (**a**) shows metaplots of ChIP-seq and ATAC-seq signals across all protein-coding genes (i, *n =* 22,258), proximal intergenic regions (ii, *n =* 35,553), and distal intergenic regions (iii, *n =* 11,418). Read density was normalized using spike-in controls added to an equal number of cells. Data represent the mean of three biological replicates for H3K9me3, H3K27me3, and H3K36me3 ChIP-seq, and two replicates for H3K4me3 and ATAC-seq, with shading indicating SEM. Significance was tested using a two-sided Kolmogorov-Smirnov test, with *P*-values indicated. **(B)** Heatmaps showing correlated changes in H3K4me3 ChIP-seq (i) and ATAC-seq (ii) signals in IAA-treated CTD-AID ESCs compared with those in early embryonic stages. Regions with significantly increased (related to totipotency peaks, *n =* 1,249/1,731, red) or decreased (related to pluripotency peaks, *n =* 4,357/7,570, blue) H3K4me3 enrichment at 24-h IAA treatments overlap with the broad H3K4me3 signals in early cleavage embryos. ATAC-seq heatmaps show regions with significantly increased accessibility (related to totipotency peaks, *n =* 971/4,113, red) and decreased accessibility (related to pluripotency peaks, *n =* 4,211/5,411, blue) around the peak center at 24 h, reflecting totipotent embryo characteristics. The numbers before/after the slash correspond to the peaks that are significantly upregulated or downregulated (after the slash) in CTD-truncated cells, and the number of overlapping peaks that are higher (totipotency peaks) or lower (pluripotency peaks) in two-cell embryos compared to ICM (before the slash). See details in Methods. **(C)** IGV snapshots showing ATAC-seq and selected histone marks in representative minor-ZGA, pluripotency, and bivalent *HOX* genes. **(D-E)** Imaging analysis revealing drastic nuclear reorganization in CTD-AID ESCs treated with IAA for 24 hours. Panel (**D**) shows DNA (i) and RNA (ii) FISH analysis of major (Maj SAT, red) and minor (Min SAT, green) satellite repeats. Panel (**E**) shows RNA FISH (i) analysis of 45S pre-rRNA (red) and 28S rRNA (green), along with immunofluorescence analysis of the nucleolus marker protein FBL (red, ii). Scale bar, 5 μm.

In CTD-truncated cells, the genome-wide gain of H3K9me3 and ATAC peaks is positively correlated, but negatively with H3K36me3 (Figures 6B, D). This suggests a chromatin state conducive to basal transcription activity. Similarly, in early cleavage embryos, paternal H3K9me3-marked genes display an open chromatin state that permits transcription due to the absence of repressive HP1α and linker histone H1 in newly established H3K9me3 chromatin before the eight-cell stage^97,98^. Intriguingly, genes with lower elongation velocity demonstrate higher ATAC signals in both control and CTD-truncated cells (Figure 6E). We postulate that the increased chromatin residency of CTD-truncated Pol II, manifested by its promiscuous binding and slow elongation, may contribute to this relatively open chromatin state. This activity may be counterbalanced by H3K9me3 deposition, potentially safeguarding genome stability.

RNA and DNA fluorescence in situ hybridization (FISH) revealed a gradual activation of major satellite repeats starting at 3 hours after IAA addition, coinciding with the breakdown of large chromocenters into smaller, weaker foci (Figures 6D and S6F, G). These changes align with a burst of pericentromeric satellite RNA expression concurrent with minor ZGA, followed by a decline that facilitates pericentromeric heterochromatin organization in early embryos^99^. Moreover, CTD-truncated cells exhibited nucleolar precursor body (NPB)-like ring structures, distinct from mature nucleoli, as indicated by staining with the nucleolar marker fibrillarin (FBL) (Figures 6E and S6G). In early embryos, these NPBs initiate ribosomal transcription only by the 8-cell stage, developing into functional nucleoli^100–104^. Consistently, mature 28S rRNA was barely detectable after 24-hour IAA treatment, while low-level rRNA transcription was observed at the NPB surface by FISH analysis of 45S precursor rRNA (Figures 6E ad S6G). Taken together, these results demonstrate that CTD truncation induces transcriptional and epigenetic reprogramming and nuclear reorganization, resembling early totipotent embryos.

### CTD length and phosphorylation in totipotency reprogramming

Lastly, we investigated the impact of CTD phosphorylation and length on genic and intergenic transcription. The Pol II CTD facilitates phosphorylation-dependent interactions with numerous RNA processing proteins^13,19,20,75^. In *S. cerevisiae*, mutations that substitute Ser2 or Ser5 with nonphosphorylatable Ala (A) or phosphomimetic Glu (E) are lethal^17^, causing pleiotropic defects including inefficient capping, splicing, mRNA instability, premature termination, and elevated cryptic intragenic initiation^8,13,18,20,22,105^. The removal of 15-18 proximal repeats is also lethal in yeast, while mammalian cells require more than 23 repeats to be removed^106,107^. In mammalian cells, a truncated CTD with 31 repeats disrupts mRNA maturation into export-competent ribonucleoprotein particles^21^. Additionally, the human CTD can replace the *D. melanogaster* CTD during fly development, indicating the conserved structural organization of Pol II across species^108^.

We engineered a series of human RPB1 (hRPB1) mutants, including substitutions of Ser5 or Ser2 or both residues to Glu (S5E, S2E, or S5S2E) in all heptapeptide repeats, and mutations with varying CTD repeat lengths (0, 5, 31, and 52 repeats) (Figure 7A). These constructs were studied in mouse ESCs with an N-terminal degron-tagged endogenous RPB1 (Pol II NTD-AID) ^109^. IAA treatments effectively degraded RPB1, abolishing Pol II transcription in NTD-AID cells. Importantly, this did not induce upregulation of minor-ZGA genes or intergenic transcription (Figures 7B, C), indicating that transcriptional inhibition alone does not explain totipotency reprogramming. In NTD-AID ESCs where endogenous Pol II was depleted, expression of wild-type hRPB1 with 52 repeats (WT or 52R) fully restored pluripotency gene programs (Figures 7B, C), emphasizing RPB1’s conservation between humans and mice.

**Figure 7.**
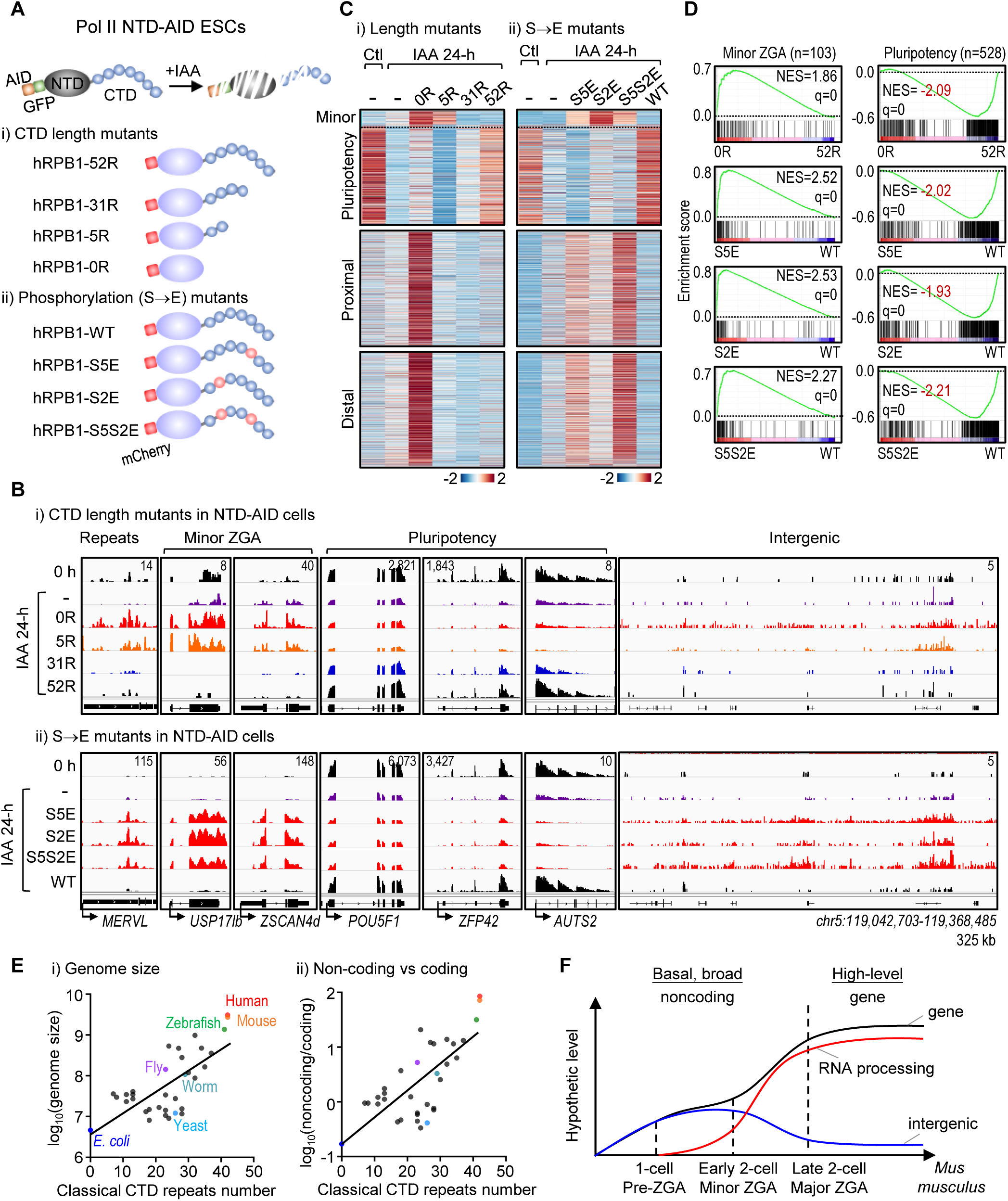
Pol II CTD length and phosphorylation in totipotent reprogramming. **(A)** Experimental scheme illustrating the expression of various human RPB1 (hRPB1) mutants in mouse Pol II NTD-AID ESCs, where the endogenous RPB1 is tagged with AID and GFP at its amino (N) terminus. The addition of IAA led to complete degradation of RPB1 in NTD-AID cells, contrasting with the carboxyl terminal truncation observed in CTD-AID cells. Panel (i) shows hRPB1 constructs with varying repeat numbers in the CTD, ranging from zero (0R) to the full-length 52 copies (52R). Panel (ii) shows hRPB1 constructs with Ser5 or Ser2 or both residues mutated to phosphomimetic Glu (S5E, S2E, or S5S2E) in all heptapeptide repeats. hRPB1-52R and hRPB1-WT serve as wild-type controls in the corresponding experiments. **(B-C)** IGV snapshots (**B**) and heatmaps (**C**) showing steady-state RNA expression in NTD-AID ESCs expressing hRPB1 with various CTD lengths (i) and phosphorylation states (ii). Various hRPB1 proteins were transiently expressed in NTD-AID ESCs treated with IAA for 24 hours, which completely degraded endogenous RPB1. The numbers of features in panels (i) and (ii) are shown before and after the slash, respectively. For genes: minor-ZGA = 97/98, pluripotency = 527/528. For intergenic bins: proximal = 33,872/34,335 and distal = 11,000/11,099. Because non-expressed features were filtered out (FPKM > 0), the numbers of features displayed in each analysis were slightly different. FPKM values and RNA read signals are the mean of two (i) or three (ii) biological replicates. **(D)** GSEA showing the upregulation of minor-ZGA genes and downregulation of pluripotency genes upon expression of hRPB1 without the entire CTD (0R) or three phosphomimetic mutants (S5E, S2E, or S5S2E) in 24-hour IAA-treated NTD-AID cells lacking the endogenous RPB1. **e,** Evolution analysis of the correlation between the number of Pol II CTD repeats with the genome size (i) and the ratio of noncoding to coding sequences (ii) across different species. Representative species are highlighted in different colors. Information on the number of CTD repeats for each species is provided in Table S10. **(F)** Schematic representation illustrating key events during zygotic gene activation (ZGA) in mouse embryos. The x-axis represents developmental stages, while the y-axis represents hypothetical levels of intergenic noncoding (blue) and protein-coding (black) transcripts, along with the degree of co-transcriptional RNA processing (red). Prior to ZGA, promiscuous and low-level transcription across the genome can be detected in mid-one-cell embryos at the PN3 stage (pre-ZGA). This early non-productive genome-wide transcription is followed by a first minor wave of transcription of a few hundred genes with fewer introns in the early two-cell stage (minor ZGA). This process may facilitate the subsequent entry of maternally stored RNA processing proteins into the nucleus. The coupling between transcription and RNA processing is then gradually established during and after major ZGA, a major wave in which thousands of protein-coding genes are highly activated.

All CTD truncation and Ser-to-Glu mutants exhibited downregulated pluripotency genes, decreased genic sense RNA signals and transcription directionality, and increased read-through (Figures 7B-D and S7A-D). Notably, hRPB1-0R, which lacks the entire CTD, and Ser-to-Glu mutants, specially the S5S2E double mutant, exhibited elevated gene-body antisense and pronounced intergenic RNA signals, along with upregulated minor-ZGA genes (Figures 7B-D and S7A, B, D). These effects mirrored those seen in CTD-AID cells and minor-ZGA embryos (Figure S7E). Intriguingly, the presence of only five repeats (5R) suppressed intergenic transcription (Figures 7B, C and S7A). Complete CTD absence or enforced phosphorylation can both increase intergenic transcription, underscoring the CTD’s crucial role and its regulated phosphorylation in restraining Pol II activity across chromatin. Expressing hRPB1 mutants with all Ser5 or Ser2 mutated to Ala (S5A, S2A, or S5S2A) had a toxic effect, inhibiting overall transcription and failing to induce reprogramming in IAA-treated NTD-AID cells (data not shown).

The hRPB1-5R mutant enhanced minor-ZGA genes without increasing intergenic transcription (Figures 7B, C and S7A), reflecting the embryonic transition during minor ZGA with limited co-transcriptional RNA processing (see Discussion). Extending the CTD to 31 repeats (31R) inhibited minor-ZGA genes and increased pluripotency gene expression, with full restoration observed with 52 repeats (52R) (Figures 7B, C). These results suggest that a longer CTD, by facilitating more efficient co-transcriptional coupling, enhances mRNA expression more effectively. Interestingly, the length of the CTD appears to correlate positively with genome size and noncoding sequences, but inversely with the proportion of protein-coding sequences across species (Figure 7E). This suggests that increasing CTD length may be an evolutionary adaptation to distinguish protein-coding mRNA production from basal ncRNA transcription, thereby addressing the challenge posed by an expanding noncoding genome in higher eukaryotes^76,77,108,110–112^.

## Discussion

Based on these results, we propose that early cleavage embryos first activate their genome through indiscriminate Pol II transcription (pre-ZGA). This is followed by the expression of a hundred short genes (minor ZGA), and ultimately, the global activation of zygotic genes (major ZGA). This transition from the genome-wide noncoding activity to protein-centered gene programs triggers development and coincides with the shift from totipotency to pluripotency and subsequent lineage differentiation (Figures 7F and S7F).

Our findings highlight the critical role of co-transcriptional RNA processing via the CTD of Pol II in driving this noncoding-to-coding transition. Regulation occurs at three levels: first, *cis*-DNA signals like introns raise RNA processing demands; second, these demands recruit numerous RBPs to the transcriptional processes; and third, the Pol II CTD coordinates transcription and RNA processing, suppressing Pol II activity in noncoding regions while promoting gene transcription. This mechanism prioritizes high-level gene expression, distinguishing the protein-coding sequences—constituting only 2% of the genome—from the prevalent low-level, short-lived, and nuclear-retained ncRNAs that dominate the genome.

Pol II mutants with a truncated CTD or carrying phosphomimetic Glu exhibited increased intergenic transcription, suggesting that the unphosphorylated CTD restricts Pol II activity on chromatin. Unlike stochastic Pol II binding at intergenic regions^113^, the enlistment of RBPs by nascent RNA to gene promoters can create local microphases that confine Pol II near promoters, a phenomenon known as pausing, which depends on the CTD^113–115^. Once a threshold is reached, these microdroplets merge into larger condensates or integrate with nearby Pol II clusters, leading to rapid CTD hyperphosphorylation^116^. This process results in longer RNA production and a burst of negative charges, altering the physicochemical properties of transcription condensates. In contrast to stochastic or enforced phosphorylation, only properly regulated CTD phosphorylation within phase-separated mRNA transcription and processing factories can effectively stabilize Pol II on chromatin for efficient elongation and RNA processing. Thus, RNA and its processing machinery regulate CTD phosphorylation to prevent stochastic Pol II behavior through phase separation and transition mechanisms.

The core RNA Pol II holoenzyme comprises 12 subunits^117,118^, while mRNA maturation involves an estimated total of over 500 proteins, with approximately 300 proteins forming spliceosomes that assemble stepwise at splicing sites to mediate RNA splicing^119–121^. Given the complexity involving hundreds of proteins, RNA processing may proceed more slowly, necessitating synchronization with transcription activity. This slower pace of mRNA maturation contrasts with the minimal processing required for ncRNA synthesis, which explains why noncoding transcription predominates before mRNA production during the awakening of zygotic genomes post-fertilization. Evolutionary analysis of exon skipping suggests transient splicing compromise during mammalian ZGA^32,122^. Pre-ZGA embryos display reduced levels of splicing regulators like SNRPB and SNRPD2, which are restored after major ZGA through increased synthesis^123^. Maternally deposited RNA packaging proteins such as hnRNP A1/C/M enter the nucleus later than RNA Pol II (30 versus 22 hphCG) ^124–127^, implying that ncRNA transcripts may facilitate their nuclear import. In minor-ZGA embryos, partial coupling with RNA processing may be sufficient for expressing shorter genes with fewer introns. It is also reported that shorter genes often undergo post-transcriptional RNA processing^128,129^. As RNA processing capacity increases, genes containing more introns and higher CpG-island density in promoters become transcribed. Transcription regulation and efficiency intensify with the need for RNA processing in genes with higher intron counts^26,27,29,60^, highlighting that co-transcriptional coupling facilitates efficient elongation in a CTD-length-dependent manner. Disrupting this coupling through CTD truncation resets genome-wide transcription, chromatin, and nuclear organization to a totipotency state, similar to pre- or minor-ZGA embryos. Apparently, there is a positive correlation between developmental potential and noncoding transcription, with totipotent embryos exhibiting higher ncRNA activity than pluripotent ICM and ICM-derived ESCs, which display more active transcription compared to multipotent stem cells and somatic cells^130,131^. This underscores the role of noncoding transcription in establishing cellular plasticity and shaping cell fates development^93,132–134^. Clarifying the distinction between protein-coding and noncoding regulation enhances our understanding of the mammalian genome, expanding the central dogma of molecular biology.

## Acknowledgments

We thank professors J. Wang at Sun Yat-Sen University, B. Li at Shanghai Jiaotong University, D. Peng and F. Tang at Peking University, Y. Yin at Zhejiang University, N. Liu and S. Luo at Tsinghua University, and the Shen Laboratory members for discussions and suggestions. This work was supported by the National Natural Science Foundation of China (32161133016 and 31925015 to X.S.); Beijing Advanced Innovation Center for Genomics; and Center for Life Sciences.

## Author contributions

X.S. supervised the project. J.X. designed and performed most experiments and bioinformatic analyses. X.L. conducted EU-staining, histone ChIP-seq and hRPB1 mutant rescuing experiments. X.Hao wrote bioinformatic scripts. X.Hu and S.M. assisted with RNA half-life and single-cell transcriptomic data analysis. D.Y. conducted mass spectrometry under H.D.’s supervision. J.Z. performed embryo microinjection under Z.C.’s supervision. X.J. provided Pol II AID ESCs. Y.H., X.J. and J.N provided suggestions. X.S. and J.X. wrote the manuscript.

## STAR METHODS

### RESOURCE AVAILABILITY

#### Lead contact

Further information and requests for resources and reagents should be directed to and will be fulfilled by the Lead Contact, Xiaohua Shen.

#### Materials availability

All unique/stable reagents generated in this study are available from the lead contact with a completed Materials Transfer Agreement.

#### Data and code availability

- Sequencing data have been deposited at GEO and are publicly available as of the date of publication. Accession numbers are listed in the key resources table.
- This paper does not report original code.
- Any additional information required to reanalyze the data reported in this paper is available from the lead contact upon request.

### EXPERIMENTAL MODEL AND SUBJECT DETAILS

#### Animal maintenance

Wild-type C57BL/6 strain mice were purchased from Tsinghua Animal Center. Mice were maintained under specific-pathogen-free conditions with a 12/12 h light/dark cycle in an environment of 20-22 °C and 55 ± 10% humidity. All animals were maintained according to the guidelines of the Institutional Animal Care and Use Committee of Tsinghua University, Beijing, China.

#### Early embryo collection

For embryo collection, C57BL/6 female mice were first injected with pregnant mare serum gonadotropin, followed 48 hours later by 5 IU of hCG (Ningbo Hormone Product Co., 110251283). Post hCG administration, the females were mated with C57BL/6 males. Embryos at the late one-cell, late two-cell, and four-cell stages were collected at 30, 48, and 60 hours following hCG administration, respectively. The collected embryos were cultured in KSOM medium (Sigma-Aldrich, MR-101-D).

#### Cell culture

The V6.5 mouse embryonic stem cell (mESC) line, including Pol II CTD-AID ESCs and Pol II NTD-AID ESCs, was derived from the inner cell mass of C57BL/6 × 129/sv crossed mice and was a gift from Ji Xiong of Peking University^78,109^. mESCs were maintained in DMEM (Thermo Fisher, 10566016), supplemented with 15% heat-inactivated fetal bovine serum (FBS, Gemini, 900-108), GlutaMAX (100× stock, Corning, 35050-061), Penicillin-Streptomycin Solution (100× stock, Corning, 30-002-CL), nonessential amino acids (100× stock, Corning, 11140-050), nucleoside mix (100× stock, Millipore, ES-008-D), sodium pyruvate (100× stock, GIBCO, 11360070), 0.1 mM 2-mercaptoethanol (Sigma, M3148), 1000 U/ml recombinant leukemia inhibitory factor (LIF), 1 μM PD0325901 (Selleck, S1036), and 3 μM CHIR99021 (Selleck, S2924). The LIF and two inhibitors (2i) were freshly added before use. The mESCs were seeded onto 0.1% gelatin (Sigma, G1890)-coated dishes and cultured in a humidified incubator at 37 °C with 5% CO_2_. All cell lines tested negative for mycoplasma.

To induce reprogramming in RPB1 AID ESCs, degron cells were pre-treated with 1 μg/ml Doxycycline hyclate (Selleck, S4163) for 12 hours to induce TIR1 expression. Subsequently, cells were treated with 500 μM auxin analog indole-3-acetic acid (IAA, Sigma, I5148) at various time points to truncate RPB1-CTD (Pol II CTD-AID ESCs) or fully degrade RPB1 (Pol II NTD-AID ESCs) via the TIR1-proteasome complex. Meanwhile, LIF and 2i were withdrawn from the ESC culture medium.

### METHOD DETAILS

#### Splicing and transcription inhibition of mouse embryos

For AMO injections, each zygote embryo was injected with 5 pl of AMO solution (1 mM). Injections were performed using an Eppendorf Transferman NK2 micromanipulator. The AMO sequences are provided in Table S11. For PlaB and DRB treatment, zygotes were obtained 22 hours post hCG injection and cultured in KSOM medium containing either DMSO, 1 μM PlaB, or 100 nM DRB for 8, 16, and 38 hours until reaching the late one-cell, late two-cell, and four-cell stages, respectively.

#### EU staining in mouse embryos

For EU staining, embryos obtained from *in vitro* fertilization were collected at different developmental stages, specifically at the middle one-cell, late one-cell, and late two-cell stages, corresponding to 10, 16, and 36 hours post-insemination, respectively. The embryos were cultured in KSOM medium with or without 100 nM DRB, supplemented with 5 mM 5-ethynyl uridine (EU; Jena Bioscience, CLK-N002), for 3 hours at 37 °C before fixation. Subsequently, the embryos were fixed with 1% paraformaldehyde (PFA; Sigma, P6148) for 12 hours at 4 °C. After fixation, the embryos were permeabilized with 0.5% Triton X-100 in PBS for 2 hours at 4 °C. The subsequent Click-iT reaction was carried out as previously described^135^. Finally, the embryos were stained with Hoechst, mounted, and imaged.

#### Plasmids and cell line construction

For the generation of human RPB1-CTD length mutants, we deleted 52 CTD repeats to create the 0 CTD repeats variant. For the 5 CTD repeats variant, we retained the 1st, 2nd, 3rd, 51st, and 52nd CTD repeats. The 31 CTD repeats variant retained the 1st to 3rd and 25th to 52nd CTD repeats. In all constructs, the non-repetitive C-terminal sequence of human RPB1 was preserved^136^. For the phosphorylation mutants of human RPB1 CTD, serine residues at positions 2 and/or 5 of all CTD repeats were substituted with alanine or glutamic acid. Additionally, hRPB1 in the phosphorylation mutant group was mutated to confer α-amanitin resistance, as described in the literature^136^.

The CAG promoter-driven human RPB1 coding sequence (CDS) with an N-terminal mCherry tag was introduced into the PiggyBac vector. This vector was co-transfected with the pBase vector into Pol II NTD-AID mESCs. Positive cells were selected using 150 μg/ml hygromycin (Yeason, 31282-04-9) for 5 days. Following selection, degron cells were treated with 1 μg/ml doxycycline hyclate for 12 hours to induce TIR1 expression and degradation of endogenous mouse RPB1 in the presence of 500 μM IAA for 24 hours. The sequences of plasmids are listed in Table S11.

#### Pol II Immunoprecipitation (IP) and mass spectrometric analysis

For mass-spectrometric analysis of both full-length and truncated RPB1, samples were prepared from four 15 cm plates of Pol II CTD-AID ESCs. Cells were treated with Dox for 12 hours and IAA for 24 hours, washed with DPBS, trypsinized, and harvested. The cell pellets were further washed with DPBS. The cell pellets were resuspended in 600 μl RIPA buffer per 15 cm plate (140 mM NaCl, 10 mM Tris, pH 8.0, 1% Triton, 0.1% sodium deoxycholate, 1 mM EDTA, 1/100 protease inhibitor (PI), and 1 μl benzonase (Sigma, E1014)). The suspension was passed through a 1 ml syringe (26 5/8 Gauge needle) 10 times and rotated at 4 °C for 30 minutes. Lysates were centrifuged at 14, 000 for 10 minutes at 4 °C.

Protein G magnetic beads (Thermo Fisher, 10003D) were washed three times with RIPA buffer and incubated with RPB1 antibody (Abclonal, A11181) at 4 °C with rotation for 1 hour (30 μl beads per 15 cm plate, 4 μl antibody per 30 μl beads). Antibody-bound beads were washed three times with RIPA buffer and incubated with cell lysates at 4 °C with rotation for 2 hours. Beads were washed three times with RIPA buffer. Finally, beads from four 15 cm plates were combined and eluted with 1× SDS buffer at 100 °C twice.

Eluted samples were loaded onto a 6% gradient SDS/PAGE gel and stained with Colloidal Blue (Thermo Fisher, LC6025). Gel bands corresponding to the intact and truncated RPB1 were reduced with DTT, alkylated with iodoacetamide, and digested with a mix of trypsin and TLCK-treated alpha chymotrypsin.

Eluted samples were loaded onto a 6% gradient SDS/PAGE gel and stained with Colloidal Blue (ThermoFisher, LC6025). Gel bands corresponding to the intact and truncated RPB1 were reduced with DTT, alkylated with iodoacetamide, and first digested with trypsin at 37℃ overnight and then digested with chymotrypsin at 25℃ overnight. Extracted peptides were analyzed by nano LC-MS/MS (UltiMate™ 3000 coupled to Fusion, Thermo Scientific), separated by reverse-phase chromatography using a gradient from 4% B/96% A to 90% B/10% A in 60 minutes (A: 0.1% formic acid, B: 0.1% formic acid in acetonitrile). MS data were queried against the CTD sequence with tryptic/ chymotryptic constraints using Byonic (version 5.0.20). Oxidation of methionine was allowed as variable modification and Carbamidomethyl of Cysteine was allowed as a fixed modification. Precursor ion mass tolerance was set at 20 ppm for all MS acquired in the Orbitrap mass analyzer and fragment ion mass tolerance was set at 0.02 Da for all MS2 spectra acquired. Matched peptides were filtered using < 1% False Discovery Rate.

#### Isolation and mass spectrometric analysis of insoluble chromatin and RNP mesh fraction

Pol II CTD-AID cells were cultured in SILAC medium containing DMEM (Thermo Fisher, 88420), supplemented with 15% FBS (Gibco, 26400-044), GlutaMAX, Penicillin-Streptomycin Solution, nonessential amino acids, Sodium Pyruvate, 0.1 mM 2-mercaptoethanol, heavy (^13^C_6_, ^15^N_2_ L-lysine-2HCl (Thermo Fisher, 88432) and ^13^C_6_, ^15^N_4_ L-arginine-HCl (Thermo Fisher, 88434)) amino acids. The LIF and two inhibitors (2i) were freshly added before use. Cells were treated with IAA for 24 hours and mixed with an equal number of untreated cells cultured in light medium. The medium was exchanged for different treatments to provide an additional biological replicate and exclude medium bias^113^.

Cells from a 15 cm plate (approximately 1.5×10^7^ cells) were trypsinized and collected in DPBS. All subsequent steps were performed under RNase-free conditions. Nuclei were isolated by resuspending the cells in 0.6 ml of cold hypotonic buffer A (20 mM Hepes, pH 7.5, 10 mM KCl, 1.5 mM MgCl_2_, 10% glycerol, 0.1% NP-40, 1 mM EDTA, 0.1 mM Na_3_VO_4_, 1 mM DTT, 1 mM PMSF, 1/100 PI). The resuspended cells were transferred to a 1 ml dounce homogenizer and homogenized with 8 strokes. Nuclei were pelleted by centrifugation at 4 °C for 5 minutes at 1430×g. The cytosolic supernatant was discarded, and permeabilized nuclei were washed in hypotonic buffer A without NP-40, followed by centrifugation for 3 minutes at 400×g. The nuclear pellet was gently resuspended in 200 μl of cold glycerol buffer (20 mM Tris, pH 7.9, 75 mM NaCl, 0.5 mM EDTA, 50% glycerol, 0.1 mM Na_3_VO_4_, 1 mM DTT, 1 mM PMSF, 1/100 PI) using wide orifice tips. An additional 200 μl of cold nuclei lysis buffer (20 mM Hepes, pH 7.6, 0.3 M NaCl, 7.5 mM MgCl_2_, 0.2 mM EDTA, 1% NP-40, 0.1 mM Na_3_VO_4_, 1 mM DTT, 1 mM PMSF, 1/100 PI) was added, followed by pulsed vortexing and incubation on ice for 1 minute. Samples were centrifuged for 2 minutes at 15,000 rpm 4 °C. Subsequently, 200 μl DPBS was added to the remaining chromatin and RNP mesh pellet, followed by brief vortexing. Samples were centrifuged for an additional 5 minutes.

The chromatin and RNP mesh and whole cell input were treated with 200 μl of RIPA buffer (25 mM Tris-HCl, pH 7.6, 280 mM NaCl, 1% NP-40, 1% sodium deoxycholate, 0.1% SDS, 1/100 PI) and subjected to sonication for 30 seconds (10 seconds on, 20 seconds off, at 25% power). Samples were centrifuged at 15,000 rpm for 10 minutes at 4 °C, and the supernatant was collected. 1 ml acetone was added to the supernatant, and proteins were precipitated overnight at -20 °C. The precipitate was centrifuged at 15,000 rpm for 30 minutes at 4 °C, washed once with cold acetone, air-dried for 5 minutes, and dissolved in 100 μl of 8 M urea in DPBS.

Protein identification was performed using the MaxQuant platform, and protein abundance was assessed by the intensity of the sum of all peptide intensities relative to the number of observable peptides of a protein (iBAQ intensity). We selected 577 proteins that consistently decreased in the chromatin and RNP mesh fraction in both replicates (IAA/Ctl < 1) while not decreasing in the input (IAA/Ctl ≥ 1) as proteins enriched on chromatin and RNP mesh fraction that decreased after 24 hours of IAA treatment. These proteins were used for gene ontology analysis.

#### Smart-seq2 library preparation and sequencing

Smart-seq2 libraries of mouse embryos were prepared following the previously described protocol^137^. Embryos were washed three times in KSOM medium and then lysed in 2 μl of lysis buffer with RNase inhibitor. Spike-in RNA (ERCC) was added according to the number of embryos. Libraries were constructed using the VAHTS TruePrepTM DNA Library PrepKit V2 for Illumina (Vazyme, TD502) and sequenced on a Novaseq 6000 platform.

#### RNA-seq and data analysis

For RNA-seq analyses of Pol II CTD-AID ESCs with time-course IAA treatment and wash-off, hRPB1-CTD length mutants, and hRPB1-CTD phosphorylation mutants in Pol II NTD-AID ESCs, purified RNAs were subjected to ribosomal RNA depletion using the Ribo-off rRNA Depletion Kit (Vazyme, N406-02). Strand-specific RNA libraries were constructed using the VAHTS Universal V6 RNA-seq Library Prep Kit (Vazyme, N604-02) and sequenced on a Novaseq 6000 or DNBSEQ-T7 platform. Drosophila S2 cells were added as spike-in controls before RNA extraction to indicate cell number.

Raw reads from RNA-seq were mapped to the mouse genome (mm10) using HISAT2 (version 2.1.0) and additionally to the Drosophila genome (dm6) using Bowtie2 (version 2.2.8) ^138^. Read counts of genes (exon) were calculated using featureCounts (version 2.0.1) ^139^ with Gencode vM25 annotation (56,262 annotated genes). Reads counts of intergenic regions were analyzed using BEDTools (version 2.27.1) ^140^, and only uniquely mapped reads were used for all analyses about intergenic region, including Fragments per kilobase of exon model per million mapped reads (FPKM) calculations, meta bin analysis, and dysregulated bins analysis. FPKM were calculated using the formula:

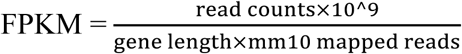

FPKM values were normalized to spike-in reads by multiplying the normalization score called F1, which was calculated by normalizing the ratio of (spike-in mapped reads/mm10 mapped reads) to the control sample as 1. All FPKM values in this study were spike-in normalized. We used averaged FPKM from two or three biological replicates for further analysis.

We identified dysregulated protein-coding genes or intergenic regions using DESeq2 (version 1.42.1), including only those genes with a total exon reads count in the analyzed samples greater than 5 (for protein-coding genes) or greater than 20 (for intergenic bins). Differentially expressed hits were identified based on log_2_(exp/Ctl) and P-value (determined using a two-sided t-test) with two or three biological replicates (see figure legends for detailed criteria). During analysis, spike-in normalization was used as input with a spike-in factor called F2 that was the ratio of (spike-in reads/Ctl-rep1 spike-in reads) to the control sample as 1. Metaplots of averaged signals in indicated regions were analyzed using ngs.plot (version 2.63) and normalized by F1. Metaplots are presented as mean ± SEM of two or three replicates. Heatmaps were drawn using pheatmap (version 1.0.12), with colors normalized by row across all analyzed samples. RNA-seq tracks were produced by bamcoverage (version 3.5.2), with --normalizeUsing 1/F2, represented as spike-in normalized signals, then merging two or three replicates using WiggleTools (version 1.2.11) and visualized by Integrative Genomics Viewer (IGV, version 2.17.0) ^141^.

#### EU-seq and data analysis

EU-seq was performed as previously described with modifications^142^. ESCs were labeled with 1 mM EU for 10 minutes. Cells were harvested in TRIzol, and spike-in control (EU labeled S2 cells) was added according to cell number. Purified RNA (5-10 μg) was treated with DNase I (RQ1, 37 °C, 30 minutes), fragmented (94 °C, 5 minutes), and ethanol precipitated. EU-labeled RNAs were used to perform the biotinylation reaction as follows: 50 mM HEPPS pH 7.5, 2.5 mM Tris(3-hydroxypropyltriazolylmethyl)amine (THPTA, Sigma, 762342), 2.5 mM CuSO_4_, 4 mM Biotin-PEG3-azide (Aladdin, B122225), 10 mM sodium ascorbate for a 1-hour reaction at room temperature. The reaction was stopped with 5 mM EDTA, and biotinylated RNA was purified by phenol-chloroform extraction. Two rounds of biotin-affinity purification were performed using M-280 Streptavidin Dynabeads (Thermo Fisher, 11205D). Eluted RNA was used for cDNA synthesis and strand-specific library construction using the Scale ssDNA-seq Lib Prep Kit (Abclonal, RK20222) and sequenced on a Novaseq 6000 or DNBSEQ-T7 platform.

Data processing was performed similarly to RNA-seq data analysis with modifications. Adapters and low-quality reads were removed, and polyC tracts were cut from the reads using Trim_galore (version 0.6.6). For identifying dysregulated genes, heatmap analysis, and GSEA analysis, FPKM was calculated using featureCounts with the parameter -t exon. For comparative analysis of antisense transcription of protein-coding genes in Figure 5, FPKM was calculated using featureCounts with the parameter -t gene.

We used the following published datasets in our analysis: EU-seq of mouse embryos from Mizuki (GSE235547); Total RNA-seq of mouse embryos from Abe (PRJEB7345); poly(A) RNA-seq of mouse embryos from Wu (GSE66390); poly(A) RNA-seq of mouse embryos from Zhang (GSE71434); scRNA-seq of mouse embryos from Xue (GSE44183); poly(A) RNA-seq of human embryos from Zou (GSE197265); scRNA-seq of human embryos from Tang (GSE36552); scRNA-seq of pig embryos from Zhai (PRJCA008416); rRNA depletion RNA-seq of zebrafish embryos from Bazzini (GSE148391). Data processing was performed similarly to RNA-seq data analysis, and all FPKM values were normalized to sequencing depth.

#### Pulse-chase EU-seq and data analysis

Pol II CTD-AID cells were cultured in 10 cm dishes until reaching 70%-80% confluence. For the IAA group, cells were pretreated with doxycycline for 12 hours and then with 500 μM IAA for 16 hours before pulse-labeling with 100 μM EU for 8 hours. The medium containing EU was removed, and cells were washed twice with DPBS to eliminate residual EU. Cells were subjected to continued treatment with 500 μM IAA during both the pulse and chase periods. Cells were harvested, counted, and lysed in TRIzol with 10% Drosophila S2 cells (pre-labeled with 100 μM EU for 24 hours) added as a spike-in according to the cell number. Labeled RNA was purified and used for constructing transcriptome sequencing libraries as described for EU-seq.

Data preprocessing and analysis were consistent with those described for EU-seq. Read counts for repeats were analyzed by featureCounts using the mm10 repeat mask GTF, and only uniquely mapped reads were included in the analysis. The annotation for lincRNAs was derived from the Gencode vM25 annotation, and read counts were calculated in the same manner as for protein-coding genes. Enhancer RNA annotation was sourced from a published paper^93^, and reads counts of these regions were calculated using BEDTools. PROMPTs were defined by initially selecting protein-coding genes longer than 1 kb, ensuring there was no overlap within a 2-kb range upstream or downstream. Subsequently, the regions located 2-kb upstream of the transcription start site (TSS) for these genes were chosen, and read counts on the antisense strand within these regions were calculated using BEDTools.

We expected the degradation of RNA transcripts would follow an exponential decay:

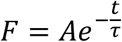

A is the amplitude parameter of degradation, and τ is the decay lifetime. So for each RNA feature, we fitted the time course data with this function in R by using minpack.lm package. Then half life of each RNA was calculated by the time of its FPKM being half of 0-h. Before fitting the function for each RNA, features with reads count = 0 in 0-h, the number of reads count = 0 in following time point are larger than 2 were filtered. Besides that, any features which were not consistent with the exponential decay curve we expected were removed.

Due to the rapid degradation of noncoding RNAs (ncRNAs) with short half-lives, their read counts become undetectable after chasing and are consequently filtered out. The ncRNA species we describe exhibit sufficient stability over a measurable period, allowing their reads to be detected. Consequently, this results in an overestimation of the average half-life range of the noncoding RNA population in the final analysis.

#### ChIP-seq and data analysis

ChIP-seq was conducted as previously described with modifications^143^. Cells were cross-linked with 1% formaldehyde for 10 minutes at room temperature and quenched with 125 mM glycine for 5 minutes. Fixed cells were washed twice with cold DPBS, counted, and human 293T cells were added as spike-in controls. Antibodies were added to Protein G-coupled Dynabeads (Thermo Fisher, 10009D) in dilution buffer and incubated with rotation for 1 hour at 4 °C. Coated beads were washed once with dilution buffer at 4 °C for 5 minutes with rotation. The antibodies used included RPB1-NTD (CST, 14958), H3K4me3 (CST, 9751), H3K36me3 (Active motif, 61021), H3K27me3 (CST, 9733), and H3K9me3 (Abclonal, A22295).

For each replicate, 1×10^6^ cells were lysed in 200 µL of nuclei lysis buffer (50 mM Tris-HCl, pH 8.1, 10 mM EDTA, 1% SDS, 1 mM PMSF, 1 mM DTT, 1/100 PI) and sonicated for 1 minute (10 seconds on, 20 seconds off, 25% power). Lysates were centrifuged at 15,000 rpm for 10 minutes at 4 °C, and the supernatant was transferred to a new tube. For immunoprecipitation, 36 µl of the supernatant was combined with 144 µl of dilution buffer (16.7 mM Tris-HCl, pH 8.1, 167 mM NaCl, 1.2 mM EDTA, 1.1% Triton X-100) containing antibody-coated beads and rotated at 4 °C for 4 hours. Beads were sequentially washed with low salt IP wash buffer (20 mM Tris-HCl, pH 8.1, 150 mM NaCl, 1 mM EDTA, 1% Triton X-100, 0.1% SDS, 0.1% Na-deoxycholate), high salt IP wash buffer (20 mM Tris-HCl, pH 8.1, 500 mM NaCl, 1 mM EDTA, 1% Triton X-100, 0.1% SDS, 0.1% Na-deoxycholate), LiCl IP wash buffer (10 mM Tris-HCl, pH 8.1, 250 mM LiCl, 1 mM EDTA, 0.5% NP-40, 0.5% Na-deoxycholate), and TE Buffer (10 mM Tris-HCl, pH 8.1, 1 mM EDTA), each for 5 minutes at 4 °C with rotation. After the final wash with 10 mM Tris-HCl, pH 7.5, samples were resuspended in 10 µl of H_2_O, and 5 µl was used for Tn5 tagmentation and library construction by the TruePrep DNA Library Prep Kit V2 (Vazyme, TD502).

For the input sample, 1 µl of the supernatant before IP was mixed with 2 µl of 1 mM MgCl_2_, 5 µl of H_2_O, and 2 µl of 1.8% Triton, and incubated at room temperature for 5 minutes. 1 μl mixture was used for Tn5 tagmentation. After tagmentation, 1 µl of TN5 stop solution (250 mM EDTA, 0.2% SDS) and 0.5 µl of Proteinase K were added, and the sample was incubated at 55 °C for 30 minutes. Input DNA was purified using DNA beads (Vazyme, N411-01) and directly used for PCR.

Raw FASTQ reads were trimmed using fastp (version 0.23.4) and mapped to the mm10 genome using Bowtie2. Spike-in reads were mapped to hg19. Unmapped reads, low-quality mapped reads, and PCR duplicates were discarded.

Metaplots of averaged signals in indicated regions were analyzed using ngs.plot and normalized by F1. Metaplots are presented as mean ± SEM of two or three replicates. For metaplots of Pol II directionality, EU-seq data from the sense strand of all genes were divided by antisense strand data. For elongation velocity, EU-seq data from the sense strand of all genes were divided by Pol II ChIP-seq data.

Peak calling was performed using MACS2 (version 2.2.7.1) ^144^, with the “--broad” option for H3K4me3, H3K36me3, H3K27me3, and H3K9me3. To identify dysregulated peaks after IAA treatment, peaks derived from Ctl and IAA groups were combined, and then read counts for both groups across all peaks were calculated using BEDTools. Differentially regulated peaks were identified using DESeq2 with a cutoff of log_2_Fc(exp/Ctl) > 1 and a *P*-value < 0.05 (two-sided t-test). Significantly upregulated peaks were defined as totipotent if log_2_Fc (1c or 2c/ICM or ESC) > 0, and significantly downregulated peaks were defined as pluripotent if log_2_Fc (1c or 2c/ICM or ESC) < 0. Coverage profiles surrounding the summit of the peaks (±2 kb for narrow peaks, ±3 kb for broad peaks) were extracted and calculated using computeMatrix (version 3.5.2). For browser visualization, replicates for each biological condition were merged into one bigwig track file with spike-in normalization.

We used the following published datasets in our analysis: Pol II ChIP-seq of mouse embryos from Liu (GSE135457); Pol II ChIP-seq of mouse embryos from Abe (GSE195840); ATAC-seq of mouse embryos from Wu (GSE66390); H3K4me3 ChIP-seq of mouse embryos from Zhang (GSE71434); H3K36me3 ChIP-seq of mouse embryos from Xu (GSE112835); H3K9me3 ChIP-seq of mouse embryos from Wang (GSE97778). Data processing was performed similarly to ChIP-seq data analysis, and all FPKM values were normalized to sequencing depth.

#### ATAC-seq and data analysis

ATAC-seq was performed using the High-Sensitivity OpenChromatin Profile Kit 2.0 (Novoprotein, N248-01A) with modifications. Nuclei were prepared following the same protocol as subcellular fractionation. Analysis of ATAC-seq data was performed similarly to ChIP-seq data analysis.

#### Defining maternal, ZGA and pluripotency genes

For mouse embryos, maternal genes, minor- and major-ZGA genes, as well as pluripotency genes, were defined based on poly(A) RNA-seq data from Zhang (GSE71434). Minor-ZGA genes were identified as those not expressed or lowly expressed in growing oocytes, GV oocytes, and MII oocytes, but upregulated in early two-cell embryos, with criteria set at FPKM > 1 in early two-cell embryos and at least a three-fold upregulation compared to oocytes. Major-ZGA genes were similarly defined but were upregulated in late two-cell embryos. Genes identified as minor-ZGA genes were excluded from the major-ZGA gene category. Pluripotency genes were defined as those upregulated in blastocyst embryos, with criteria of FPKM > 1 and at least a three-fold upregulation compared to zygote, early two-cell, late two-cell, four-cell, and eight-cell embryos. Genes identified as major-ZGA genes were excluded from the pluripotency gene category. Maternal genes were defined as those expressed in MII oocytes (FPKM > 1) but that had become downregulated (by at least threefold) at the late two-cell stage. Genes identified as pluripotency genes were excluded from the maternal gene category.

For human embryos, maternal genes, minor- and major-ZGA genes, as well as pluripotency genes, were defined based on poly(A) RNA-seq data from Zou (GSE197265). Minor-ZGA genes were defined as those upregulated in four-cell embryos, with criteria of FPKM > 1 in four-cell embryos and at least a three-fold upregulation compared to MII oocytes and zygotes. Major-ZGA genes were defined as those upregulated in eight-cell embryos, with criteria of FPKM > 1 in eight-cell embryos and at least a three-fold upregulation compared to MII oocytes and zygotes. Genes identified as minor-ZGA genes were excluded from the major-ZGA gene category. Pluripotency genes were defined as those upregulated in hESCs, with criteria of FPKM > 1 and at least a three-fold upregulation compared to zygote, two-cell, four-cell, and eight-cell embryos. Genes identified as major-ZGA genes were excluded from the pluripotency gene category. Maternal genes were defined as those expressed in MII oocytes (FPKM > 1) but that had become downregulated (by at least threefold) at the eight-cell stage. Genes identified as pluripotency genes were excluded from the maternal gene category. mESC bivalent genes were previosly defined by Gao Shaorong group^96^.

For pig embryos, maternal genes, minor- and major-ZGA genes, as well as pluripotency genes, were defined based on single-cell RNA-seq from Zhai (PRJCA008416). Minor-ZGA genes were defined as those upregulated in two-cell embryos, with criteria of FPKM > 1 in two-cell embryos and at least a three-fold upregulation compared to MII oocytes. Major-ZGA genes were defined as those upregulated in four-cell embryos, with criteria of FPKM > 1 in four-cell embryos and at least a three-fold upregulation compared to MII oocytes. Genes identified as minor-ZGA genes were excluded from the major-ZGA gene category. Pluripotency genes were defined as those upregulated in blastocyst, with criteria of FPKM > 1 and at least a three-fold upregulation compared to two-cell, four-cell, and eight-cell embryos. Genes identified as major-ZGA genes were excluded from the pluripotency gene category. Maternal genes were defined as those expressed in MII oocytes (FPKM > 1) but that had become downregulated (by at least threefold) at the four-cell stage. Genes identified as pluripotency genes were excluded from the maternal gene category.

For zebrafish embryos, maternal genes, minor- and major-ZGA genes were defined based on rRNA depletion RNA-seq from Bazzini (GSE148391). Minor-ZGA genes were defined as those upregulated in 4 hours post-fertilization embryos, with criteria of FPKM > 1 and at least a three-fold upregulation compared to 0 hours post-fertilization embryos. Major-ZGA genes were defined as those upregulated in 5 hours post-fertilization embryos, with criteria of FPKM > 1 and at least a three-fold upregulation compared to 0 hours post-fertilization embryos. Genes identified as minor-ZGA genes were excluded from the major-ZGA gene category. Maternal genes were defined as those expressed in 0 hours post-fertilization embryos (FPKM > 1) but that had become downregulated (by at least threefold) at 5 hours post-fertilization embryos. Genes identified as major-ZGA genes were excluded from the maternal gene category.

#### Defining intergenic regions

Gene annotation files were downloaded from the GENCODE database, and we chose the release M25 (GRCm38.p6) of the mouse genome for analysis. All gene transcripts, containing 200 annotated lncRNAs, 5,629 lincRNAs, and 21,859 protein-coding genes (both exons and introns) are defined as genic regions. These transcripts are merged into 24,155 genic bins with a median length of 16 kb, covering 42.5% of the genome. Proximal intergenic bins were defined as regions 10 kb upstream and downstream of genic bins, resulting in 35,553 bins with a median length of 10 kb, covering 11.5% of the genome. The remaining regions were defined as distal intergenic bins, resulting in 11,418 bins with a median length of 35 kb, covering 46% of the genome.

For humans, we used the release hg38 (GRCh38.p6) of human genome annotation in the research. Genic bins were defined as regions containing annotated lncRNAs, lincRNAs, and protein-coding genes, resulting in 22,730 genic bins with a median length of 25 kb, covering 56.5% of the genome. Proximal intergenic bins were defined as regions 10 kb upstream and downstream of genic bins. These transcripts are merged into 32,655 bins with a median length of 10 kb, covering 8.9% of the genome. The remaining regions were defined as distal intergenic bins, resulting in 9,948 bins with a median length of 35 kb, covering 34.6% of the genome.

For pigs, we used the release Sscrofa11.1 of pig genome annotation in the research. Genic bins were defined as regions containing annotated lncRNAs, lincRNAs, and protein-coding genes, resulting in 27,696 genic bins with a median length of 8 kb, covering 46.8% of the genome. Proximal intergenic bins were defined as regions 10 kb upstream and downstream of genic bins. These transcripts are merged into 39,163 bins with a median length of 10 kb, covering 13.9% of the genome. The remaining regions were defined as distal intergenic bins, resulting in 11,492 bins with a median length of 33 kb, covering 39.3% of the genome.

#### Gene Set Enrichment Analysis (GSEA) and Gene Ontology (GO) analysis

Gene set enrichment analysis (GSEA) was performed using GSEA (version 2.2.4) as previously described^145^. We compared the expression of defined gene sets between control and experimental groups, including mouse minor-ZGA, major-ZGA, and pluripotency genes. All annotated genes (*n =* 56,262) in each RNA-seq dataset were used for GSEA analysis.

Gene ontology (GO) analysis was performed using the DAVID Functional Annotation Bioinformatics Microarray Analysis tool (https://david-d.ncifcrf.gov/)^146^.

#### Unsupervised hierarchical clustering analysis

The sva (version 3.52.0) R package was used to remove batch effects between data obtained from different studies. Unsupervised hierarchical clustering analysis was performed using pheatmap (version 1.0.12). Gene expression levels of replicates were merged by taking the mean. RNA-seq data generated in this study used spike-in normalization with rRNA-depletion RNA-seq data.

#### Calculation of Pol II behavior

Initially, we filtered 13,272 protein-coding genes with a length greater than 1 kb and no other gene annotations within 2-kb upstream or downstream. Gene regions were defined as follows: PROMPTs (2-kb upstream of TSS on the antisense strand), Promoter Region (-300 bp to +300 bp relative to the TSS), Gene Body (+300 bp downstream of the TSS to -300 bp upstream of the TES), and TES Downstream (2-kb downstream of the TES). For histone modification analysis, the promoter region was defined as ±500 bp around the TSS, and the gene body was defined as the region excluding the 500 bp downstream of the TSS. Read counts for Pol II ChIP data and RNA-seq data in these regions were calculated using BEDTools, with strand-specific parameters for strand-specific libraries. FPKM values were also normalized by spike-in.

Pol II behavior was quantified using the following metrics: Pausing Index (PI), calculated as the ratio of FPKM of Pol II-ChIP in the promoter region to the FPKM in the gene body; Elongation Velocity, determined by comparing the FPKM of EU-seq in the gene body to the FPKM of Pol II-ChIP in the gene body; Read-through Index, calculated as the ratio of the FPKM of EU-seq downstream of the TES to the FPKM in the gene body; Directionality of TSS, assessed by comparing the FPKM on the sense strand of the gene to the FPKM on the PROMPTs using EU-seq data; and Directionality of Gene Body, calculated by comparing the FPKM of the sense strand to the antisense strand of the gene using EU-seq data.

The calculation of Pol II behavior in human RPB1 CTD mutations was performed using rRNA-depletion RNA-seq. Results for early embryonic Pol II behavior were derived using Pol II ChIP-seq of mouse embryos from Liu (GSE135457) and Abe (GSE195840), and EU-seq of mouse embryos from Mizuki (GSE235547).

#### Immunofluorescence

For immunofluorescence, cells were fixed in 4% PFA for 15 minutes at room temperature and washed twice with DPBS. Cells were blocked and permeabilized with blocking buffer (5% BSA, 0.3% Triton X-100 in DPBS) for 1 hour at room temperature. Primary antibodies [EXOSC10 (1:1,000, Santa Cruz, 374595), FBL (1:1,000, Abclonal, A1136)] were diluted in staining buffer (1% BSA, 0.3% Triton X-100 in DPBS) and incubated for 1 hour at room temperature. Cells were washed three times with DPBS for 5 minutes each. Secondary antibodies (1:1,000 diluted in staining buffer, ABflo® 594-conjugated Goat anti-Mouse IgG, Abclonal, AS054; ABflo® 647-conjugated Goat Anti-Rabbit IgG, Abclonal, AS075) were incubated with the cells for 1 hour at room temperature, followed by three washes with DPBS for 5 minutes each. Cells were then stained with DAPI for 10 minutes at room temperature, mounted, and imaged.

#### DNA FISH for repeats sequence detection

DNA FISH for satellite repeats in mESCs was performed as previously described^147^. mESCs growing on 35 mm glass-bottom dishes were fixed with 4% PFA in DPBS for 10 minutes at room temperature, followed by a DPBS wash for 2 minutes. Permeabilization was carried out with 0.5% Triton X-100 in DPBS. After three DPBS washes, cells were treated with 0.1 M HCl for 5 minutes and then incubated with 0.1 mg/ml RNase A in DPBS for 45 minutes at 37 °C. Cells were washed three times with 2×SSC buffer before prehybridization, which was performed by incubating cells in 50% formamide in 2×SSCT (2×SSC + 0.1% Tween 20) for 5 minutes at room temperature, then for 20 minutes at 47 °C. Samples were denatured for 2.5 minutes at 78 °C on a water-immersed heat block.

For hybridization, 200 μl of hybridization buffer containing 2×SSC, 50% formamide, 20% dextran sulfate, and 0.5 μM each of the major and minor satellite probes was applied to the samples, which were then incubated in a humidified chamber at 37 °C for about 16 hours. Finally, mESCs were washed twice with 2×SSCT for 15 minutes at 50 °C, and twice with 2×SSCT for 1 hour at room temperature. Labeled mESCs were briefly rinsed with 2×SSC and mounted with DAPI.

#### RNA FISH experiments for major and minor satellite repeats

ESCs were plated in confocal dishes with Matrigel-coated coverslips at least 12 hours before the RNA FISH experiment. Cells were fixed in 4% PFA for 10 minutes at room temperature. Permeabilization was performed with 100% cold methanol for 10 minutes, followed by rehydration with 70% ethanol for another 10 minutes. The cells were then treated with 1 M Tris-HCl (pH 8.0) for 5 minutes. Hybridization was performed by covering the cells with hybridization buffer containing 0.4 μM each of the major and minor satellite probes in 100 μl 2×SSC supplemented with 1 mg/ml yeast tRNA, 0.005% BSA, 10% dextran sulfate, and 25% formamide. The cells were incubated at 37 °C for 1 hour. Post-hybridization, the dishes were washed sequentially with 4×SSC and 2×SSC at 37 °C on a shaker at 500 rpm for 30 minutes each. Cells were then stained with DAPI in 2×SSC supplemented with 0.1% Triton X-100 for 10 minutes at room temperature and mounted in Fluoromount-G.

#### smiFISH for 5’ ETS and 28S rRNA detection

For smiFISH (single molecule inexpensive FISH), primary probe sets were designed using the Probe Designer tool provided by LGC company (https://www.biosearchtech.com/stellaris-designer) (Table S11). Hybridization probes were prepared according to a previously described method^148^. Briefly, primary probes were hybridized with a fluorescently-labeled oligonucleotide (Cy3 for 5’ ETS, Cy5 for 28S) called FLAP in the following mixture: 2 μl of 20 μM primary probes, 0.5 μl of 100 μM FLAP, 2 μl of 10× NEB 3.0 buffer, and 5.5 μl of H_2_O. The mixture was incubated in a PCR cycler at 85 °C for 3 minutes, 65 °C for 3 minutes, and 25 °C for 5 minutes.

Cells were fixed with 4% PFA for 10 minutes, followed by two washes with DPBS for 5 minutes each. Next, the cells were treated with 70% ethanol for 1 hour and washed with DPBS for 5 minutes. Cells were then rinsed with wash buffer (15% formamide, 1×SSC) for 15 minutes. Hybridization was carried out by covering the slides with a drop of hybridization buffer containing 80 nM of the prepared probes (for co-staining of 5’ ETS and 28S RNA), along with 10% ethylene carbonate, 10% dextran sulfate, 2×SSC, and 0.8 μg/μl yeast tRNA. The slides were sealed and incubated at 37 °C for 1 hour. Post-hybridization, slides were washed twice with wash buffer at 40 °C on a shaker at 500 rpm for 30 minutes each. Cells were then stained with DAPI in DPBS buffer at room temperature for 10 minutes, mounted, and imaged.

#### Quantification of microscopic data

Images were captured using a Nikon A1R MP-Multiphoton microscope, Nikon A1 HD25 laser scanning confocal microscope, or Nikon AXR NSPARC super-resolution laser scanning confocal microscope. Image analysis was conducted using NIS-Elements AR Analysis software and Fiji (ImageJ) software^149^. For satellite analysis, each nucleus was identified using DAPI-stained DNA signals. Pixels above a threshold for the satellite signal were considered “satellite foci”. We quantified the number, summed area, and intensity of satellite foci per nucleus.

#### CTD number and genome complexity analysis

All Pol II CTD repeat numbers were obtained from a previously published study^77^. CTD sequences with lethal substitutions in yeast or lacking repeat characteristics were excluded. Only CTD regions that conformed to essential functional units in yeast and conserved heptapeptides without appropriate adjacent sequences to form a functional unit were counted.

Genomic information for each species was sourced from Ensembl. The CDS proportion was calculated as the length of all annotated protein-coding genes’ CDS in each species. The remaining regions were classified as noncoding, including UTRs, introns of protein-coding genes, and intergenic regions.

### QUANTIFICATION AND STATISTICAL ANALYSIS

All experiments were conducted with a minimum of two biological replicates unless otherwise specified. Statistical analyses were performed using GraphPad Prism 10.0 software, R studio (version 4.3.2), or Excel 2019. The tests used and the corresponding *P*-values are indicated in the figures and figure legends. Details of the statistical tests are provided within the figures and figure legends. Significance levels were reported as follows: > 0.1234 (ns), < 0.0332 (*), < 0.0021 (**), < 0.0002 (***), < 0.0001 (****).

