## Supplemental Figure for "Co-transcriptional RNA processing boosts zygotic gene activation"

### SUPPLEMENTARY INFORMATION

Figure S1. Inhibition of RNA splicing blocks zygotic gene activation, related to Figure 1

Figure S2. Truncation of the Pol II CTD downregulates global gene transcription but upregulates minor-ZGA genes, related to Figure 2

Figure S3. Pol II CTD truncation induces pervasive transcription across the genome resembling early embryos before major ZGA, related to Figure 3

Figure S4. Compromised RNA decay upon Pol II CTD truncation, related to Figure 4

Figure S5. The Pol II CTD enhances transcription regulation and efficiency in complex genes, related to Figure 5

Figure S6. CTD truncation induces epigenetic reprogramming and nuclear organization resembling totipotent embryos, related to Figure 6

Figure S7. Pol II CTD length and phosphorylation in totipotent reprogramming, related to Figure 7

Figure S1. Inhibition of RNA splicing blocks zygotic gene activation, related to Figure 1.

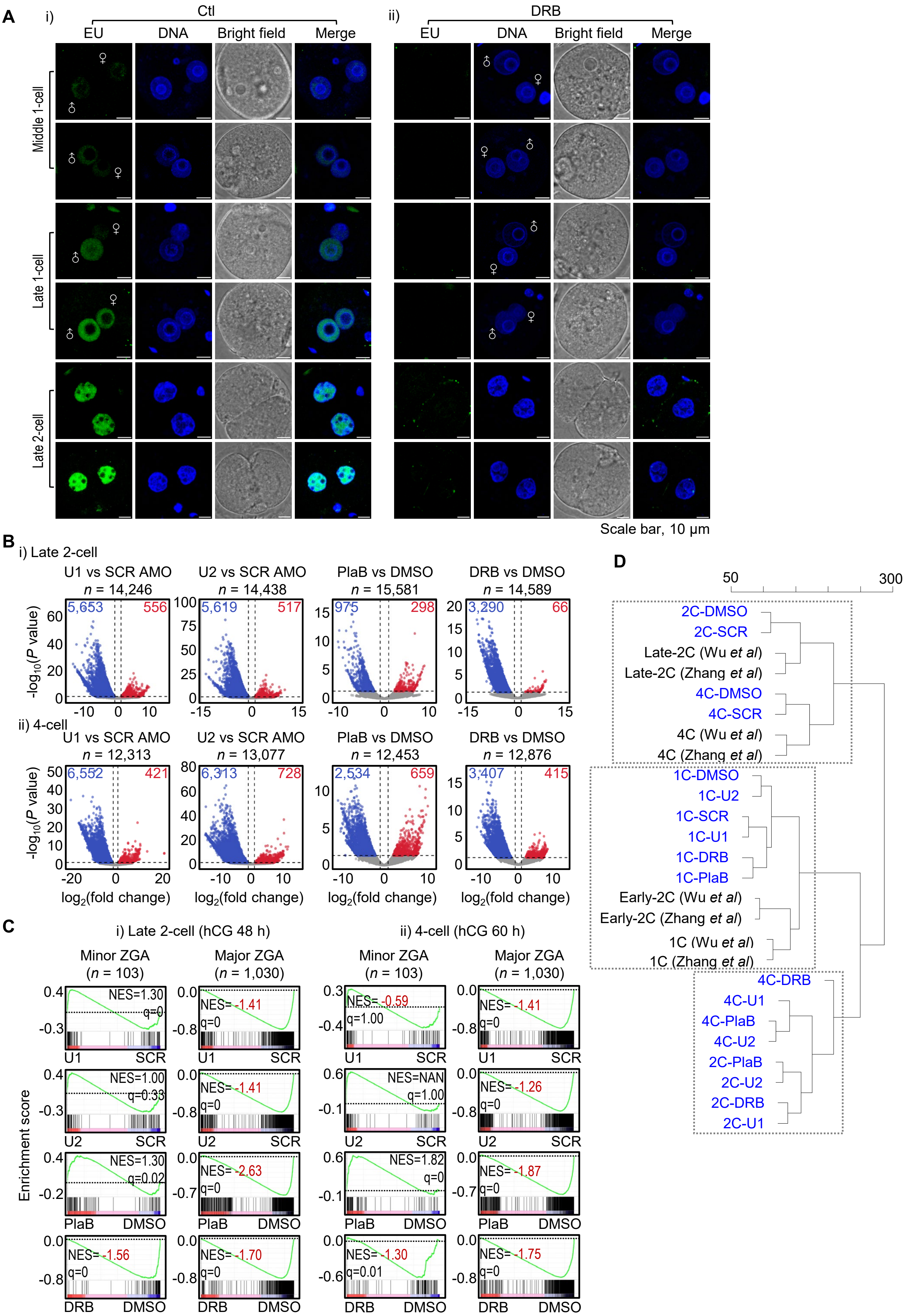

**Figure S1. Inhibition of RNA splicing blocks zygotic gene activation, related to Figure 1**

**(A)** EU staining (green) of control and DRB-treated one-cell (control, $n$ = 29; DRB, $n$ = 33) and late two-cell embryos (control, $n$ = 30; DRB, $n$ = 17) are presented. Two representative images from two independent experiments are displayed. Scale bar: 10 μm.

**(B)** Volcano plots illustrating differential gene expression in embryos treated with splicing or transcription inhibition at the late two-cell (i, 48 hphCG) and four-cell (ii, 60 hphCG) stages compared to control embryos. Only protein-coding genes with a total exon read counts > 5 in all samples were included in each individual analysis with the number of genes analyzed labeled above each plot. Genes with a $\log_2$(fold change) > 1 and $P$-values < 0.05 are highlighted in blue (downregulated) and red (upregulated), with the number of such genes annotated above each plot. Spike-in normalization was used (see details in Methods). $P$-values were obtained using a two-tailed t-test based on three biological replicates.

**(C)** GSEA of minor and major ZGA genes in embryos treated with splicing or transcription inhibition at the late two-cell (i) and four-cell stages (ii). Gene counts, normalized enrichment scores (NES), and false discovery rate (FDR) q-values are as indicated.

**(D)** Hierarchical clustering analysis of mouse embryos treated with or without splicing and transcription inhibition, based on the global transcriptome of protein-coding genes ($n$ = 22,258). The scale represents the height of clustering among samples. Both exon and intron signals were used for this analysis. Two sets of early embryo RNA-seq data are also shown here.

Figure S2. Truncation of the Pol II CTD downregulates global gene transcription but upregulates minor-ZGA genes, related to Figure 2.

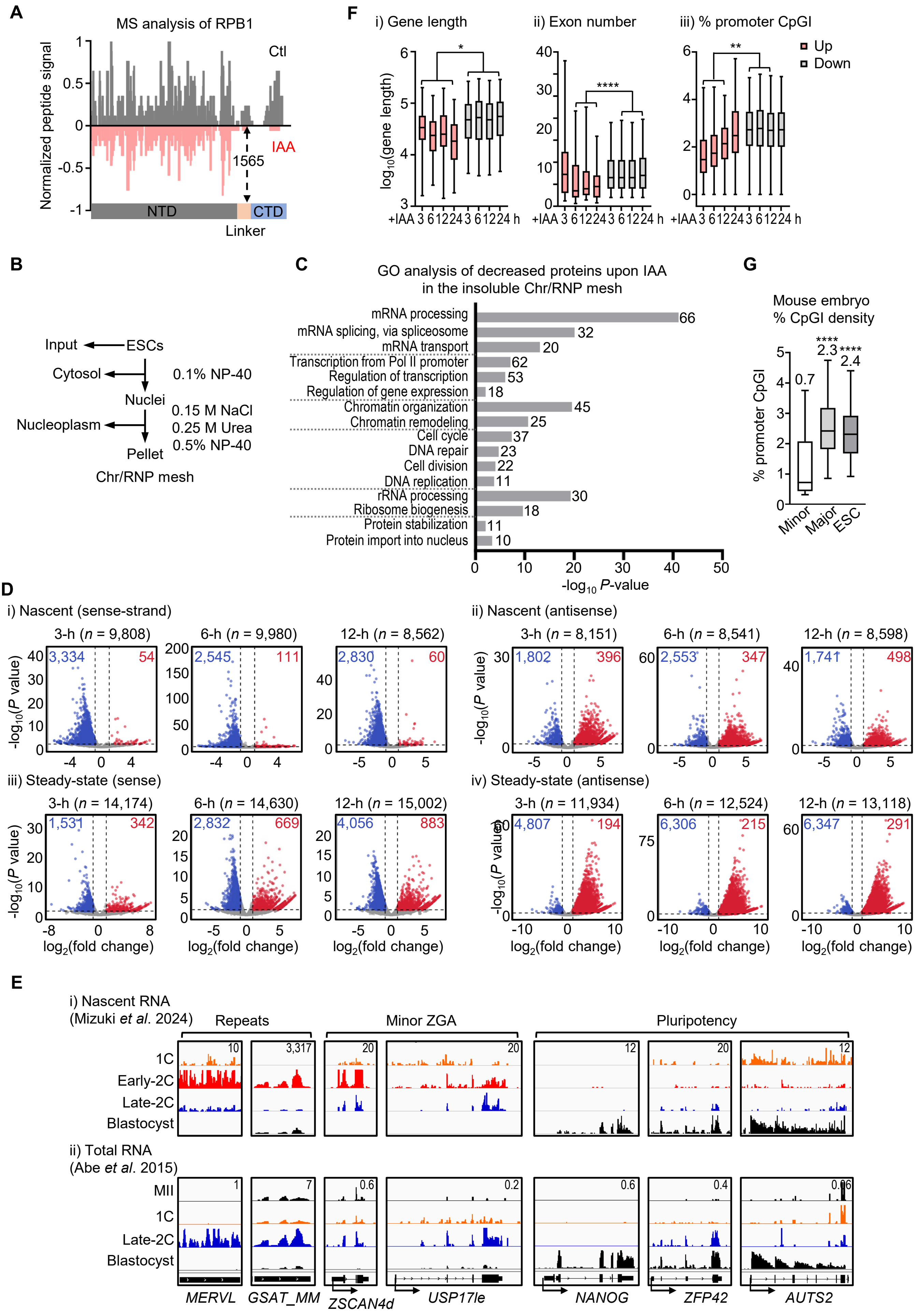

**Figure S2. Truncation of the Pol II CTD downregulates global gene transcription but upregulates minor-ZGA genes, related to Figure 2**

**(A)** Mirror plot showing summed extracted areas of peptides from full-length (Pol II CTD-AID 0 h, black) and truncated RPB1 (Pol II CTD-AID 24 h, red), normalized to the signal at residues 365-382.
**(B)** Workflow for subcellular fractionation and chromatin and ribonucleoprotein (RNP) mesh extraction followed by mass spectrometry.
**(C)** Gene Ontology (GO) analysis of proteins ($n$ = 577) significantly decreased in chromatin and RNP mesh fractions (fold change < 1, $n$ = two replicates) but not in whole-cell extracts (fold change ≥ 1, $n$ = two replicates), with the number of proteins per term labeled.
**(D)** Volcano plots show differential gene expression of sense (left) and antisense (right) strands in Pol II CTD-AID ESCs post 3, 6, and 12-hour IAA treatment compared to untreated control (0-hour) in (i) EU-seq and (ii) rRNA-depletion RNA-seq. Only protein-coding genes sense strands with a total exon read count > 5 and antisense strands with a total read count > 20 were included. The number of genes analyzed is labeled above each plot. Genes with $\log_2$(fold change) > 1 and $P$ < 0.05 are highlighted in blue (downregulated) and red (upregulated), with the number of such genes annotated above each plot. Normalization was done using spike-in controls added to an equal number of cells. $P$-values were obtained using a two-tailed t-test based on two biological replicates.
**(E)** IGV snapshots of EU-seq (i, Mizuki *et al.*) and total RNA-seq (ii, Abe *et al.*) for selected genomic loci in mouse embryos at specific stages as indicated. Transcription direction (bottom) and the signal scale (top right) are provided. The signals are represented averaging from two biological replicates for EU-seq.
**(F)** Box plots illustrate that upregulated genes are significantly shorter in length and have fewer exons and lower CpG-island density at their promoters compared to downregulated genes. This analysis was based on nascent gene expression (EU-seq) at various time points after adding IAA. The numbers of analyzed genes in the gene-length (i), exon-number (ii), and promoter CpG density (iii) analyses are shown sequentially before and after the slash, respectively. Upregulated: 3-h = 54/48/42, 6-h = 111/104/96, 12-h = 60/57/49, 24-h = 96/91/84. Downregulated: 3-h = 3,334/3,172/3,141, 6-h = 2,545/2,432/2,410, 12-h = 2,830/2,681/2,656, 24-h = 2,552/2,430/2,407. The box plots display the 5th, 25th, 50th, 75th, and 95th percentiles, excluding outliers. Statistical significance was determined using a two-tailed Mann-Whitney-Wilcoxon test, with $P$-values indicated as follows: > 0.1234 (ns), < 0.0332 (*), < 0.0021 (**), < 0.0002 (***), < 0.0001 (****). For further details, refer to Table S4.
**(G)** The statistical distribution of CpG-island density in promoters among minor-ZGA genes ($n$ = 60), major ZGA genes ($n$ = 855), and ESC-specific expressed genes ($n$ = 438) is presented as box plots. These box plots illustrate the 5th, 25th, 50th, 75th, and 95th percentiles, with the median value labeled. Statistical significance was determined using a two-tailed Mann-Whitney-Wilcoxon test, with $P$-values indicated as follows: > 0.1234 (ns), < 0.0332 (*), < 0.0021 (**), < 0.0002 (***), < 0.0001 (****).

FigureS3. Pol II CTD truncation induces pervasive transcription across the genome resembling early embryos before major ZGA, related to Figure 3.

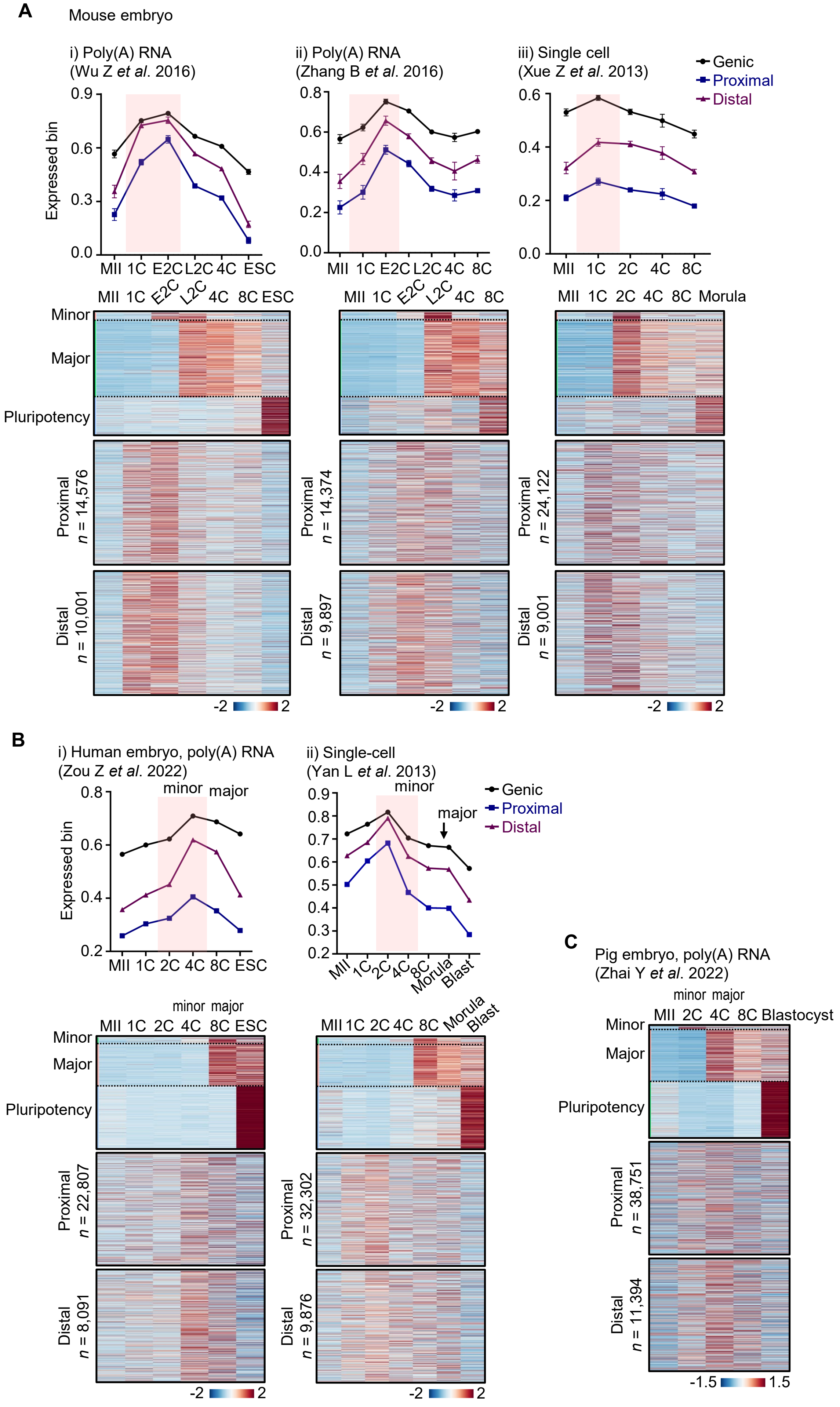


Continue to the next page

Figure S3. Pol II CTD truncation induces pervasive transcription across the genome resembling early embryos before major ZGA, related to Figure 3.

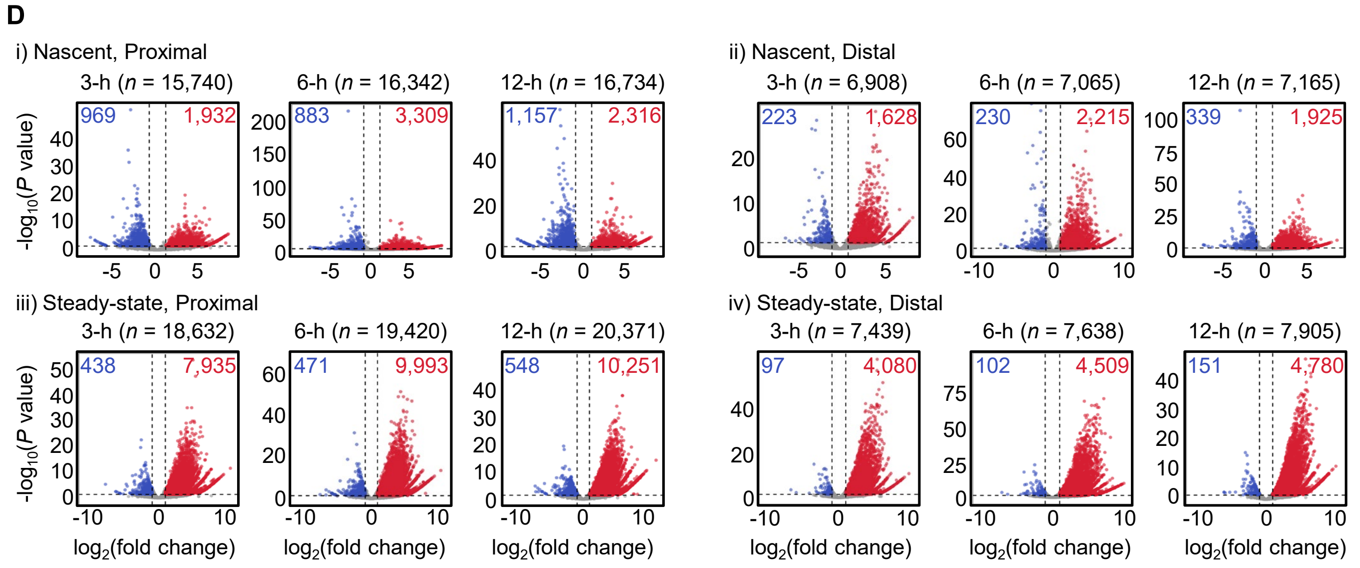


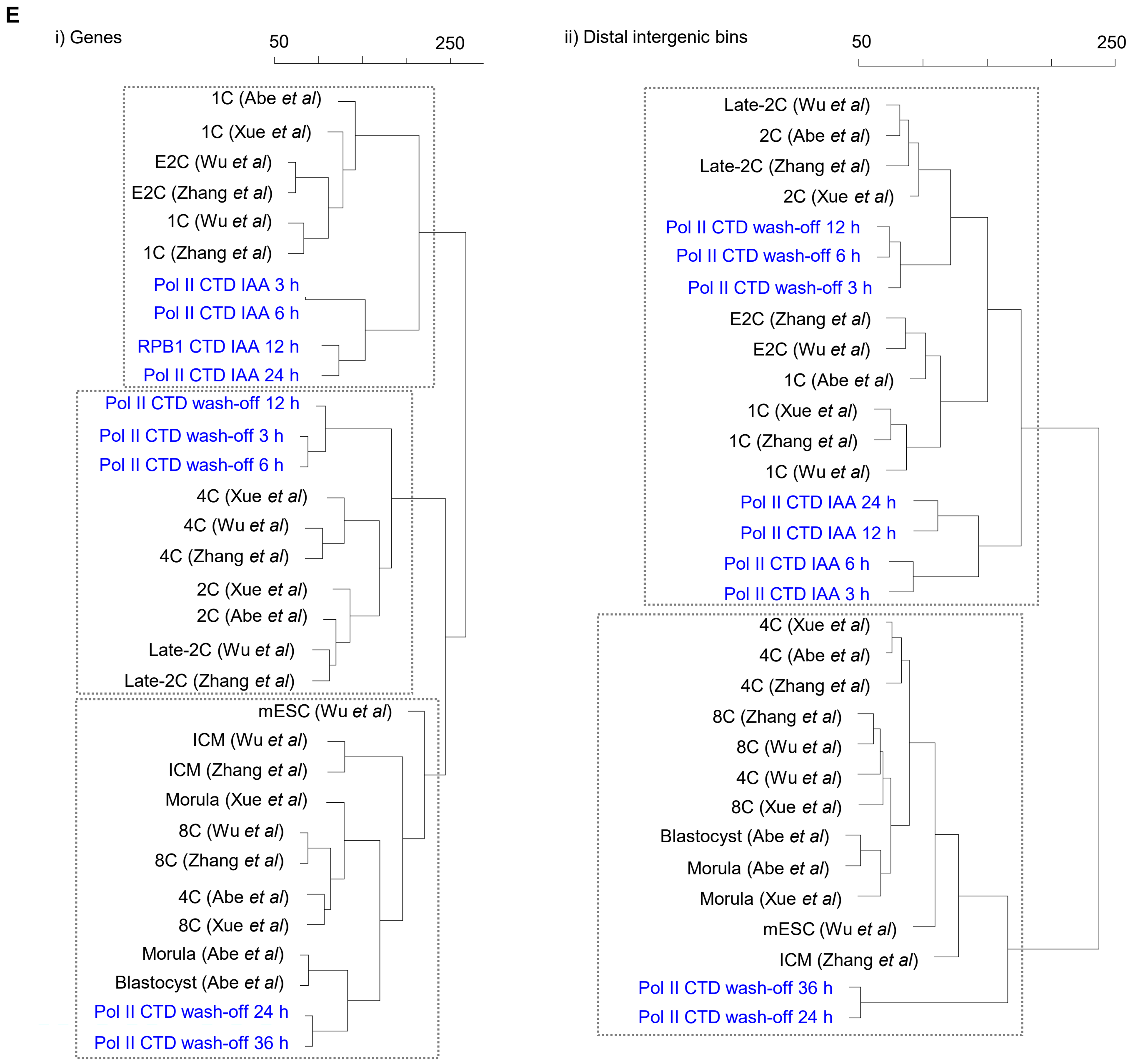

**Figure S3. Pol II CTD truncation induces pervasive transcription across the genome resembling early embryos before major ZGA, related to Figure 3**

**(A)** Analysis of steady-state RNA in early mouse embryos based on published RNA-seq. The upper panel shows the percentages of transcribed bins (FPKM > 0) during early embryonic development. The pre- and minor ZGA stages, highlighted in pink shading, exhibit the highest level of expressed genic and intergenic bins in steady-state RNA levels. This suggests a greater proportion of indiscriminate genome-wide transcription compared to other embryonic stages. The bottom heatmap illustrates the shift from noncoding to coding transcription during the transition from minor to major ZGA. Numbers of analyzed genes: 103 minor-ZGA genes, 1,030 major-ZGA genes, and 528 pluripotency genes. The FPKM values represent the mean of two biological replicates for poly(A) RNA-seq (i, ii) and three biological replicates in single-cell RNA-seq (iii), with a filtering threshold of minimum FPKM > 0.

**(B)** Analysis of steady-state RNA in early human embryos based on published RNA-seq. The upper panel shows the percentages of transcribed bins (FPKM > 0) during early embryonic development. The pre- and minor ZGA stages, highlighted in pink shading, exhibit the highest level of expressed genic and intergenic bins in steady-state RNA levels. This suggests a greater proportion of indiscriminate genome-wide transcription compared to other embryonic stages. The bottom heatmap illustrates the shift from noncoding to coding transcription during the transition from minor to major ZGA. Numbers of analyzed genes: 132 minor-ZGA genes, 914 major-ZGA genes, and 1,341 pluripotency genes, with a filtering threshold of minimum FPKM > 0.

**(C)** Heatmap illustrating the shift from noncoding to coding transcription during the transition from minor to major ZGA in pig embryos. Numbers of analyzed genes: 74 minor-ZGA genes, 1,228 major-ZGA genes, and 1,331 pluripotency genes. The FPKM values represent the mean of two biological replicates for poly(A) RNA-seq, with a filtering threshold of minimum FPKM > 0.

**(D)**Volcano plots illustrating differential expression of intergenic bins in Pol II CTD-AID ESCs post 3, 6, and 12-hour IAA treatment compared to untreated control (0-hour) in (i) EU-seq and (ii) rRNA-depletion RNA-seq. Only intergenic bins with a total read count > 20 in each individual analysis were included. The number of bins analyzed is indicated above each plot. Bins with $log_2$(fold change) > 1 and $P < 0.05$ are highlighted in blue (downregulated) and red (upregulated), with the number of such bins annotated above each plot. Normalization was done using spike-in controls added to an equal number of cells. *P*-values were calculated using a two-tailed t-test from two biological replicates.

**(E)** Hierarchical clustering analysis of Pol II CTD-AID mESCs upon IAA treatment and wash-off with four published mouse embryonic transcriptomes, based on the global transcriptome of protein-coding genes (i, $n$ = 22,258; both exon and intron signals were used for this analysis) and distal intergenic bins (ii, $n$ = 11,418). The scale represents the height of clustering among samples.

Figure S4. Compromised RNA decay upon Pol II CTD truncation, related to Figure 4.

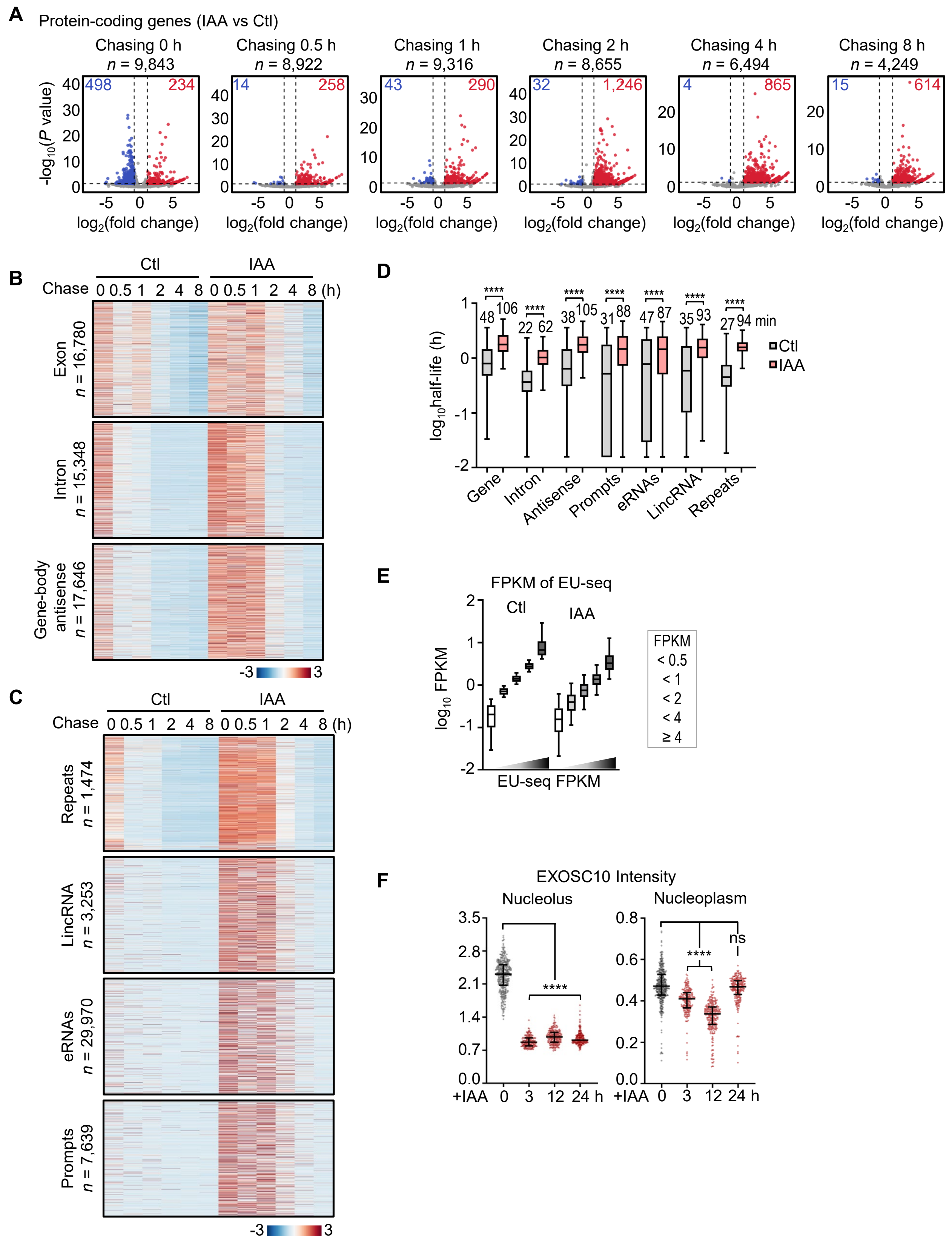

**Figure S4. Compromised RNA decay upon Pol II CTD truncation, related to Figure 4**

**(A)** Volcano plots illustrating the differential expression of protein-coding genes in Pol II CTD-AID ESCs treated with IAA compared to untreated controls during a chase period of 0-8 h. Only protein-coding genes with a total exon read count greater than 5 in each individual analysis were included. The number of genes analyzed is indicated above each plot. Genes with $\log_2$(fold change) > 1 and *P* < 0.05 are highlighted in blue (downregulated) and red (upregulated), with the number of such genes annotated above each plot. Normalization was performed using spike-in controls added to an equal number of cells. *P*-values were calculated using a two-tailed t-test from two biological replicates.
**(B)** Heatmap showing EU-labeled RNA abundance of all mRNAs (*n* = 16,780), introns (*n* = 15,348), and antisense transcripts (*n* = 17,646) of protein-coding genes at different time points during the chase. The '*n*' values denote the number of intergenic bins with maximum FPKM > 0. Expression levels were averaged from two biological replicates using spike-in normalization and z-score normalized.
**(C)** Heatmap showing EU-labeled RNA abundance of all repeats (*n* = 1,474), lincRNAs (*n* = 3,253), enhancer RNAs (*n* = 22,970), and promoters (upstream 2-kb of gene TSS, *n* = 7,639) at different time points during the chase. The '*n*' values denote the number of RNA with maximum FPKM > 0. Expression levels were averaged from two biological replicates using spike-in normalization.
**(D)** Box plots showing the half-life of RNA transcripts from protein-coding genes (including exons and introns, *n* = 13,154), gene introns (*n* = 11,846), antisense transcripts of protein-coding genes (*n* = 12,343), PROMPTs (upstream antisense 2-kb of gene TSS, *n* = 2,027), eRNAs (*n* = 8,045), lincRNAs (*n* = 1,122), and repeats (*n* = 1,282) with and without IAA treatment in Pol II CTD-AID ESCs. Only genes or regions with detectable RNA reads at 30 minutes or beyond can be used for half-life calculation, so fast-decaying transcripts within the 30-minute chase cannot be computed, potentially overestimating RNA half-lives; however, all analyzed transcripts show enhanced stability. The box plots display the 5th, 25th, 50th, 75th, and 95th percentiles, with median values labeled. Statistical significance was determined using a two-sided paired Mann-Whitney-Wilcoxon test, with *P*-values < 0.0001 (****).
**(E)** Categorization of genes based on different expression levels in Pol II CTD-AID ESCs (without IAA treatment) using the average FPKM of two biological replicates from EU-seq: FPKM < 0.5 (*n* = 1,634), < 1 (*n* = 1,392), < 2 (*n* = 1,702), < 4 (*n* = 1,296), and ≥ 4 (*n* = 1,086). After 24 hours of IAA treatment, the expression levels of genes in different categories decreased, but the relative ranking among categories remained unchanged.
**(F)** Immunofluorescence analysis of EXOSC10 in Pol II CTD-AID ESCs with IAA addition for 0-24 hours. Quantification of the EXOSC10 signal in the nucleolus and nucleoplasm is shown. Number of cells (*n*) analyzed in the nucleolus: 0 h, *n* = 372; 3 h, *n* = 182; 12 h, *n* = 226; 24 h, *n* = 227. Number of cells (n) analyzed in the nucleoplasm: 0 h, *n* = 409; 3 h, *n* = 203; 12 h, *n* = 270; 24 h, *n* = 241. The scatter dot plots display the 25th, 50th, and 75th percentiles. Statistical significance was determined using a two-sided paired Mann-Whitney-Wilcoxon test, with *P*-values denoted as follows: > 0.1234 (ns), < 0.0332 (*), < 0.0021 (**), < 0.0002 (***), < 0.0001 (****).

Figure S5. The Pol II CTD enhances transcription regulation and efficiency in complex genes, related to Figure 5.

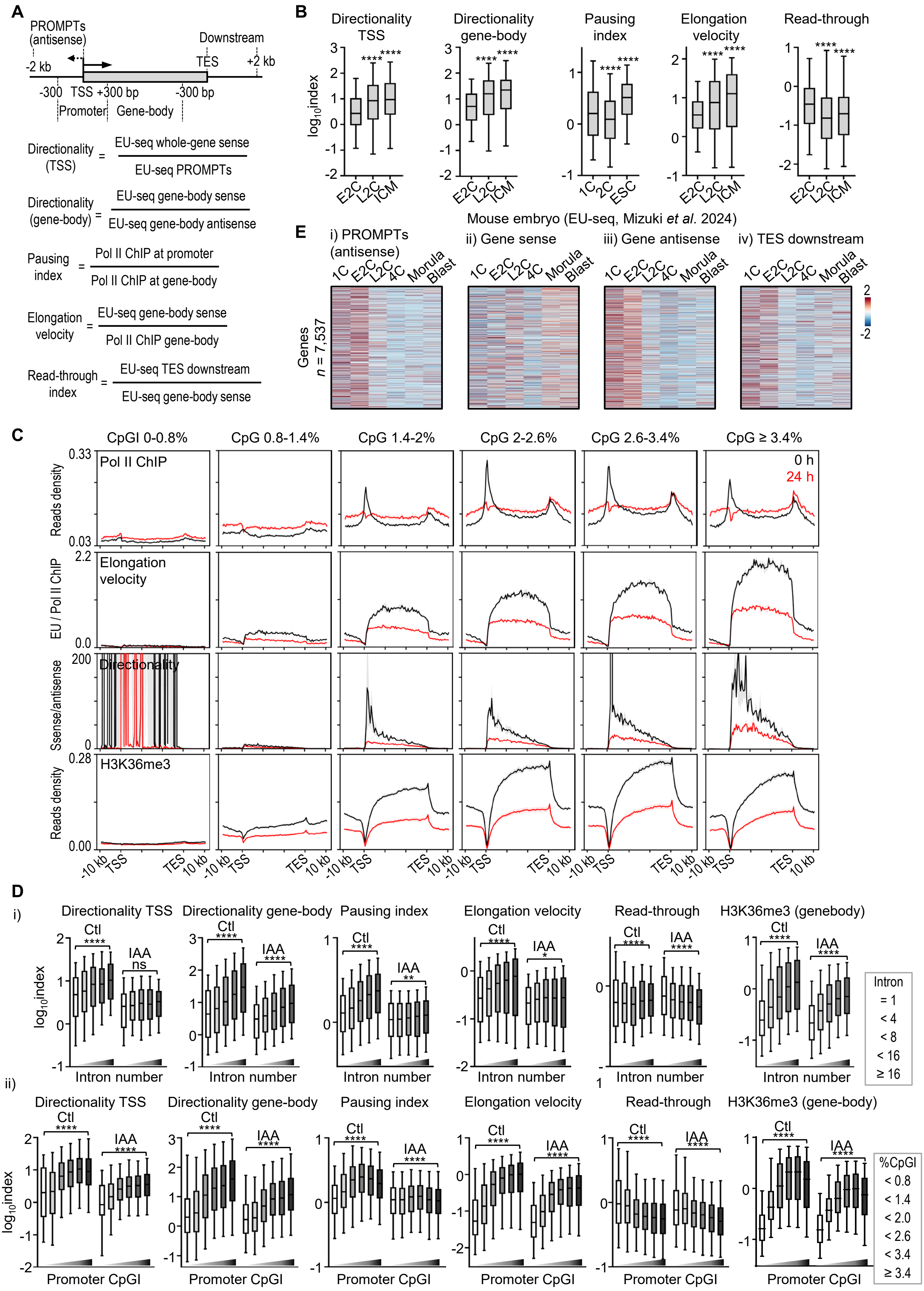

**Figure S5. The Pol II CTD enhances transcription regulation and efficiency in complex genes, related to Figure 5**

**(A)** Diagram illustrating the definitions for various genomic regions, including PROMPTs (gene TSS upstream 2-kb on the antisense strand), promoter (gene TSS ± 300 bp), gene body (gene TSS downstream 300 bp to TES upstream 300 bp), and gene TES-downstream region (gene TES-downstream 2-kb). Formulas used to calculate transcription directionality at TSS, directionality within gene bodies, pausing index, elongation velocity, and read-through index for Figure 5d and Extended Data Fig. 5b, d are also shown.
**(B)** Box plots illustrating the decreases in Pol II directionality around TSS ($n$ = 1,401) and within gene bodies ($n$ = 10,404), promoter pausing index ($n$ = 3,228), and elongation velocity ($n$ = 9,055), as well as an increase in read-through transcription beyond TES ($n$ = 2,435) in mouse embryos at the one-cell or early two-cell (minor ZGA) compared with late two-cell (major ZGA), and blastocyst (post-ZGA) stages. Specific calculation methods are detailed in Extended Data Fig. 5a and Methods. The box plots display the 5th, 25th, 50th, 75th, and 95th percentiles, with outliers omitted. Statistical significance was determined using a two-tailed Mann-Whitney-Wilcoxon test, with $P$-values denoted as follows: > 0.1234 (ns), < 0.0332 (*), < 0.0021 (**), < 0.0002 (***), < 0.0001 (****). Mouse embryo EU-seq (Mizuki *et al.*) and Pol II ChIP-seq (Liu *et al.* and Kenichiro *et al.*) data were used for these calculations.
**(C)** Metaplots showing Pol II binding, directionality, elongation velocity, and the elongation histone mark H3K36me3 at distinct classes of genes categorized by promoter CpG-island density before and after a 24-hour IAA treatment.
**(D)** Box plots showing the correlation between exon number (i) or CpG-island density of the gene promoter (ii) with Pol II behavior and the elongation histone mark H3K36me3 in Pol II CTD-AID ESCs upon IAA treatment at 0 h and 24 h. Exon number categories in directionality of TSS/gene body/pausing index/elongation velocity/read-through/H3K36me3: = 2 ($n$ = 121/278/694/375/203/1,250), < 4 ($n$ = 408/1,089/1,112/1,067/606/2,018), < 8 ($n$ = 814/2,687/1,757/2,181/1,270/3,046), < 16 ($n$ = 903/3,707/1,667/2,639/1,522/3,038), ≥ 16 ($n$ = 639/2,797/1,010/1,902/1,033/1,980). CpG-island density categories in directionality of TSS/gene body/pausing index/elongation velocity/read-through index/H3K36me3: < 0.8% ($n$ = 45/362/1,415/605/120/2,445), < 1.4% ($n$ = 219/1,127/896/1,125/433/1,872), < 2.0% ($n$ = 520/1,901/1,052/1,518/913/1,855), < 2.6% ($n$ = 667/2,343/1,084/1,580/1,062/1,742), < 3.4% ($n$ = 668/2,290/936/1,469/1,006/1,593), ≥ 3.4% ($n$ = 660/1,800/738/1,428/920/1,604). The box plots display the 5th, 25th, 50th, 75th, and 95th percentiles, with outliers omitted. $P$-values were calculated using an unpaired, one-way ANOVA Kruskal-Wallis test, with $P$-values denoted as follows: > 0.1234 (ns), < 0.0332 (*), < 0.0021 (**), < 0.0002 (***), < 0.0001 (****).
**(E)** Heatmaps displaying nascent transcription levels (mouse embryo EU-seq, Mizuki *et al.*) across various genomic regions. These include PROMPTs (2-kb upstream of gene TSS on the antisense strand), gene sense (from TSS to TES on the sense strand), gene antisense (from TSS to TES on the antisense strand), and TES-downstream regions (2-kb downstream of TES on the sense strand) for 7,537 genes showing detectable signals (maximum FPKM > 0) in each region. Each row represents different regions of the same gene. Expression levels were averaged from two biological replicates.

Figure S6. CTD truncation induces epigenetic reprogramming and nuclear organization resembling totipotent embryos, related to Figure 6.

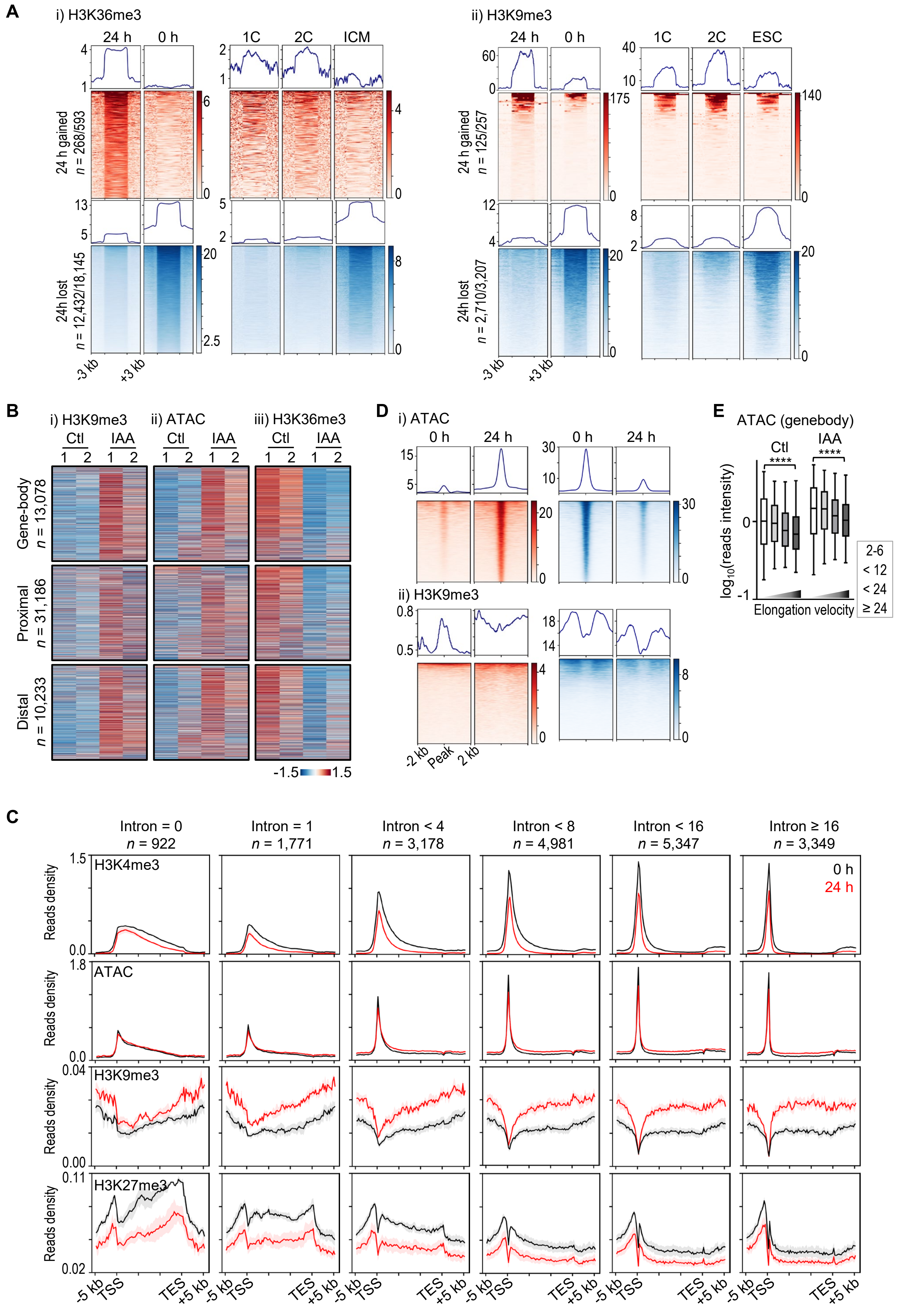


Continue to the next page

Figure S6. CTD truncation induces epigenetic reprogramming and nuclear organization resembling totipotent embryos, related to Figure 6.

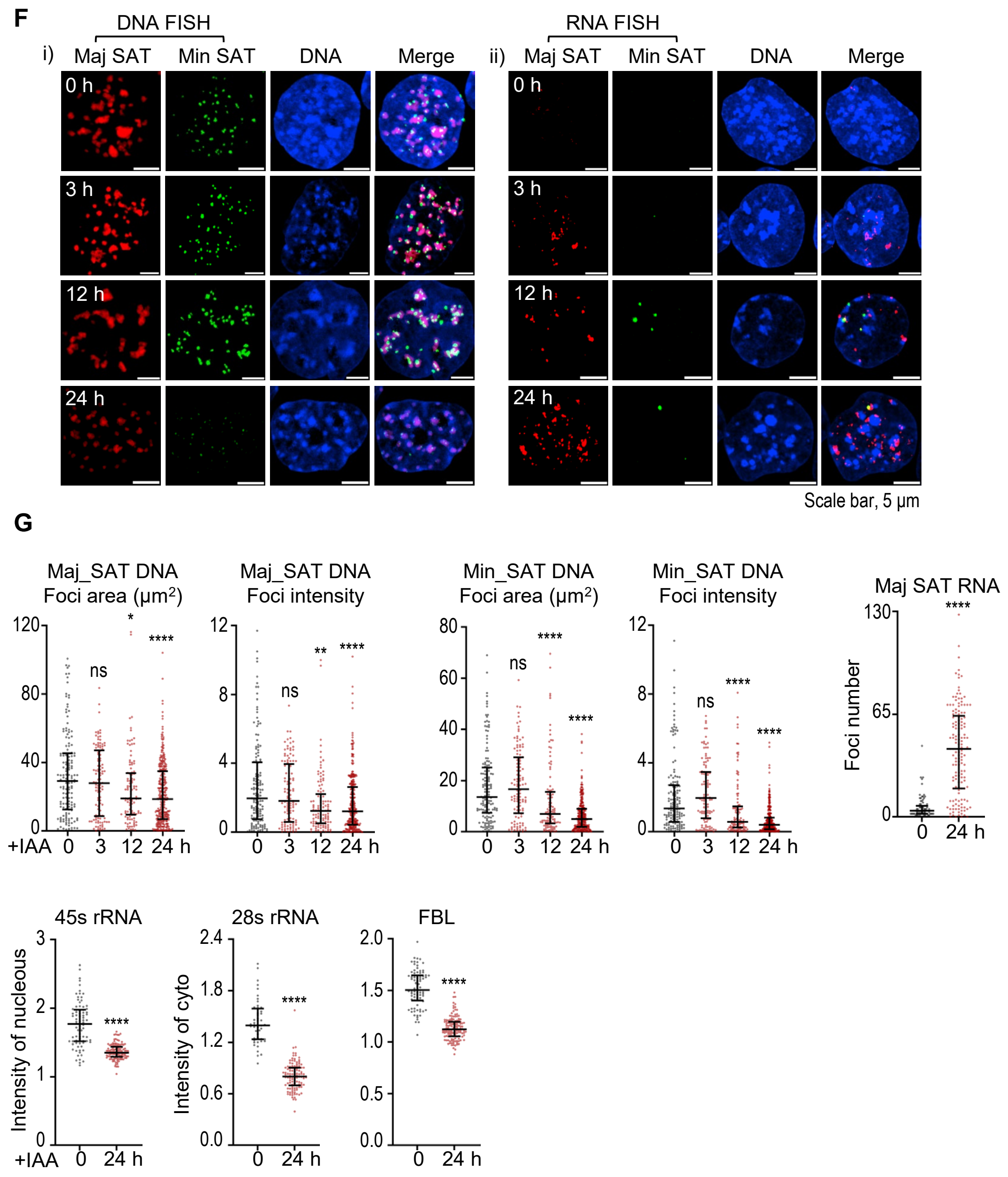

**Figure S6. CTD truncation induces epigenetic reprogramming and nuclear organization resembling totipotent embryos, related to Figure 6**

**(A)** Heatmaps of H3K36me3 (i) and H3K9me3 (ii) ChIP-seq signals in Pol II CTD-AID ESCs at 0 h and 24 h post-IAA treatment, compared with early embryonic stages. H3K36me3 heatmaps highlight regions with significantly increased (*n* = 268/593, red) and decreased (*n* = 12,432/18,145, blue) enrichment at 24h, associated with totipotent embryo features. H3K9me3 heatmaps show regions with significantly increased (*n* = 125/257, red) and decreased (*n* = 2,710/3,207, blue) enrichment at 24h, associated with totipotent embryo features. The numbers before and after the slash correspond to the peaks that are significantly upregulated or downregulated (after the slash) in CTD-truncated cells, and the number of overlapping peaks that are higher (totipotency peaks) or lower (pluripotency peaks) in two-cell embryos compared to ICM/ESC (after the slash). See details in Methods.
**(B)** Heatmaps showing H3K9me3 enrichment, chromatin openness, and H3K36me3 enrichment for all protein-coding genes (*n* = 13,078), proximal intergenic regions (*n* = 13,078), and distal intergenic regions (*n* = 13,078). Each row represents the same gene or bin across the three heatmaps with a filtering threshold of maximum FPKM > 0.
**(C)** Metaplots showing H3K4me3, ATAC, H3K9me3 and H3K27me3 at distinct classes of genes categorized by intron number before and after a 24-hour IAA treatment.
**(D)** Heatmaps of ATAC-seq (i) and H3K9me3 (ii) ChIP-seq signals in regions with significantly increased accessibility (*n* = 4,114, red) and decreased accessibility (*n* = 5,413, blue) around the peak center in Pol II CTD-AID ESCs at 24 h post-IAA treatment.
**(E)** Box plots showing the correlation between Pol II elongation velocity and ATAC-seq signals in gene bodies at 0 h and 24 h post-IAA treatment. Elongation velocity categories: < 2 (*n* = 3,566), < 6 (*n* = 1,919), < 12 (*n* = 1,613), < 24 (*n* = 1,815), ≥ 24 (*n* = 905). The box plots display the 5th, 25th, 50th, 75th, and 95th percentiles, with outliers omitted. *P*-values were calculated using an unpaired, one-way ANOVA Kruskal-Wallis test, with *P*-values < 0.0001 (****).
**(F)** DNA FISH (i) analysis of major (Maj DNA, red) and minor (Min DNA, green) satellite repeats post-IAA treatment for 3-24 hours, alongside RNA FISH (ii) analysis of the same repeats at identical time points. Scale bar, 5 μm.
**(G)** Quantification of immunofluorescence and FISH signals in Pol II CTD-AID ESCs upon IAA treatment for 0-24 hours. Number of cells (n) analyzed for major satellite DNA: 0 h, *n* = 147; 3 h, *n* = 114; 12 h, *n* = 109; 24 h, *n* = 309. Number of cells (*n*) analyzed for minor satellite DNA: 0 h, *n* = 148; 3 h, *n* = 112; 12 h, *n* = 107; 24 h, *n* = 307. Number of cells (*n*) analyzed for major satellite RNA: 0 h, *n* = 83; 24 h, *n* = 151. Number of cells (n) analyzed for 45S rRNA: 0 h, *n* = 82; 24 h, *n* = 115. Number of cells (*n*) analyzed for 28S rRNA: 0 h, *n* = 48; 24 h, *n* = 111. Number of cells (*n*) analyzed for FBL: 0 h, *n* = 82; 24 h, *n* = 136. The scatter dot plots depict the 25th, 50th, and 75th percentiles, with statistical significance determined via a two-sided paired Mann-Whitney-Wilcoxon test, and *P*-values indicated as follows: > 0.1234 (ns), < 0.0332 (*), < 0.0021 (**), < 0.0002 (***), < 0.0001 (****).

Figure S7. Pol II CTD length and phosphorylation in totipotent reprogramming, related to Figure 7.

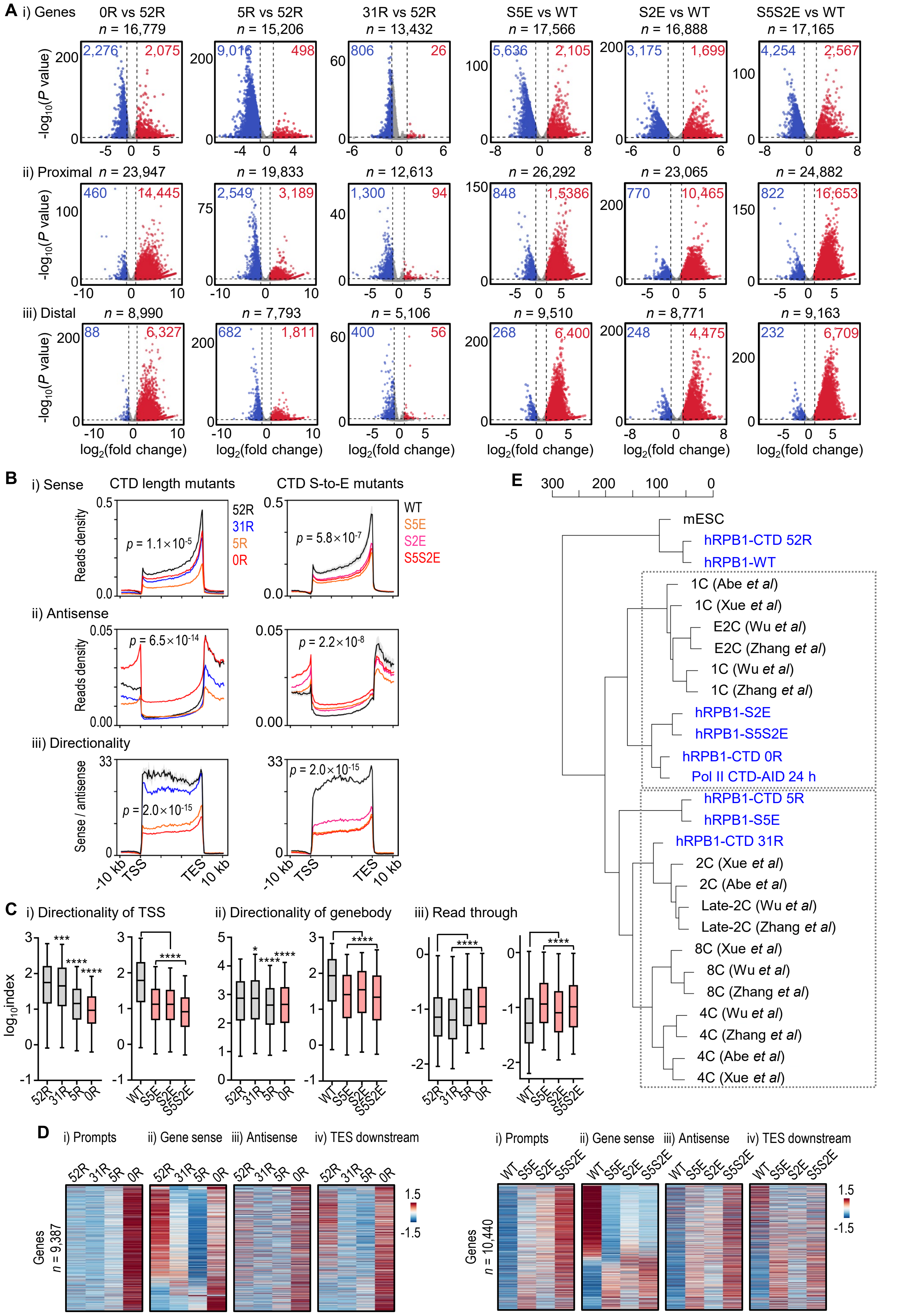


Continue to the next page

Figure S7. Pol II CTD length and phosphorylation in totipotent reprogramming, related to Figure 7.

**F**

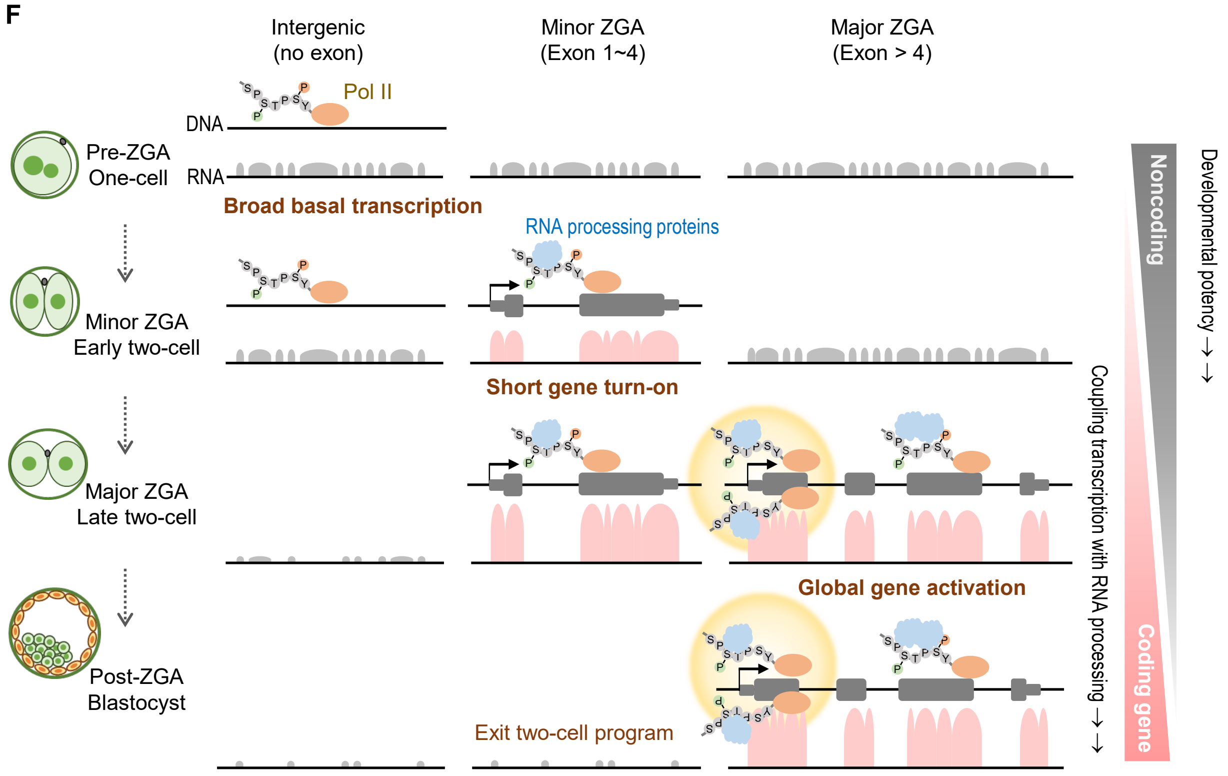

**Figure S7. Pol II CTD length and phosphorylation in totipotent reprogramming, related to Figure 7**

**(A)** Volcano plots show differential gene and intergenic bin expression in Pol II NTD-AID mESCs upon IAA treatment for 24 h while expressing human RPB1 mutants with different CTD repeat numbers and phosphorylation abilities compared to wild-type hRPB1 in rRNA-depletion RNA-seq. Only protein-coding genes with a total exon read counts > 5 and intergenic bins with a total read count > 20 were included. The number of genes and bins analyzed is labeled above each plot. Genes and bins with a $\log_2$(fold change) > 1 and $P < 0.05$ are highlighted in blue (downregulated) and red (upregulated), with the number of such genes and bins annotated above each plot. Normalization was done using spike-in controls added to an equal number of cells. *P*-values were obtained using a two-tailed t-test based on two biological replicates for CTD length mutants and three replicates for CTD phosphorylation mutants.

**(B)** Metagene analysis of hRPB1-CTD length (left) and hRPB1-CTD phosphorylation (right) mutants in Pol II NTD-AID ESCs after the addition of IAA treatment for 24 hours. RNA read signals in the sense (i), antisense (ii) strand and directionality (iii) of all protein-coding genes ($n$ = 22,258) are shown. Read density was normalized to cell numbers using spike-in controls. Data represent mean values from two biological replicates for hRPB1-CTD length and three replicates for hRPB1-CTD phosphorylation, with shadings representing SEM. Significance was tested with a two-sided Kolmogorov-Smirnov test, with *P*-values as indicated. In the CTD length mutants, the P-value corresponds to the significance of the difference between 0R and 52R.

**(C)** Box plots illustrating the decreases in Pol II directionality around TSS ($n$ = 3,963/6,214) and within gene bodies ($n$ = 13,026/15,371), as well as an increase in read-through transcription beyond TES ($n$ = 6,103/7,194) in hRPB1-CTD length/phosphorylation mutants. Specific calculation methods are detailed in Extended Data Fig. 5a and Methods. The box plots display the 5th, 25th, 50th, 75th, and 95th percentiles, with outliers omitted. Statistical significance was determined using a two-tailed Mann-Whitney-Wilcoxon test, with *P*-values denoted as follows: > 0.1234 (ns), < 0.0332 (*), < 0.0021 (**), < 0.0002 (***), < 0.0001 (****).

**(D)** Heatmaps showing transcription levels in CTD length (i) and phosphorylation (ii) mutants across various genomic regions. These include PROMPTs (2-kb upstream of gene TSS on the antisense strand), gene sense (from TSS to TES on the sense strand), gene antisense (from TSS to TES on the antisense strand), and TES-downstream regions (2-kb downstream of TES on the sense strand) for protein-coding genes showing detectable signals (maximum FPKM > 0) in each region. Each row represents different regions of the same gene.

**(E)** Hierarchical clustering analysis of hRPB1 mutants in Pol II NTD-AID mESCs with four published mouse embryonic transcriptomes, based on the global transcriptome of protein-coding genes (i, $n$ = 22,258; both exon and intron signals were used for this analysis). The scale represents the height of clustering among samples.

**(F)** Cartoon illustration of our model: we propose that early cleavage embryos first activate their genome through indiscriminate Pol II transcription (pre-ZGA). This is followed by the expression of a hundred short genes (minor ZGA), and ultimately, the global activation of zygotic genes (major ZGA). This shift from the genome-wide noncoding activity to the initiation of protein-centered gene programs triggers development, coinciding with the transition from totipotency to pluripotency and lineage differentiation. Our findings demonstrate that co-transcriptional RNA processing via the CTD of Pol II plays a key role in driving the noncoding-to-coding transition.
